# Structural control for the coordinated assembly into functional pathogenic type-3 secretion systems

**DOI:** 10.1101/714097

**Authors:** Nikolaus Goessweiner-Mohr, Vadim Kotov, Matthias J. Brunner, Julia Mayr, Jiri Wald, Lucas Kuhlen, Sean Miletic, Oliver Vesper, Wolfgang Lugmayr, Samuel Wagner, Frank DiMaio, Susan Lea, Thomas C. Marlovits

## Abstract

Functional injectisomes of the type-3 secretion system assemble into highly defined and stoichiometric bacterial molecular machines essential for infecting human and other eukaryotic cells. However, the mechanism that governs the regulated step-wise assembly process from the nucleation-phase, to ring-assembly, and the filamentous phase into a membrane embedded needle complex is unclear. We here report that the formation of a megadalton-sized needle complexes from *Salmonella enterica* serovar Typhimurium (SPI-1, *Salmonella* pathogenicity island-1) with proper stoichiometries is highly structurally controlled competing against the self-assembly propensity of injectisome components, leading to a highly unusual structurally-pleiotropic phenotype. The structure of the entire needle complex from pathogenic injectisomes was solved by cryo electron microscopy, focused refinements (2.5-4 Å) and co-variation analysis revealing an overall asymmetric arrangement containing cyclic, helical, and asymmetric sub-structures. The centrally located export apparatus assembles into a conical, pseudo-helical structure and provides a structural template that guides the formation of a 24-mer cyclic, surrounding ring, which then serves as a docking interface comprising three different conformations for sixteen N-terminal InvG subunits of the outer secretin ring. Unexpectedly, the secretin ring excludes the 16^th^ protein chain at the C-terminal outer ring, resulting in a pleiotropic 16/15-mer ring and consequently to an overall 24:16/15 basal body structure. Finally, we report how the transition from the pseudo-helical export apparatus into the helical filament is structurally resolved to generate the protein secretion channel, which provides the structural basis to restrict access of unfolded effector substrates. These results highlight the diverse molecular signatures required for a highly coordinated assembly process and provide the molecular basis for understanding triggering and transport of unfolded proteins through injectisomes.

## Introduction

Molecular machines ^1, 2^ can only perform their function in the cell if they assemble properly ^3^. Hence, mis-assembly into non-functional or defective systems is energetically costly and could even result in unwanted side-effects to the cell. As a consequence, assembly processes of complex molecular machines composed of one or many copies of numerous components are tightly regulated including various control elements at the transcriptional, translational, post-translational or structural level. These elaborate controls ensure a spatiotemporal confined and defined order of assembly steps into functional complexes, such as the ribosome or the spliceosome, executed in a crowded cellular environment ^4, 5^.

Type-3 secretion systems (T3SSs) assemble at bacterial membranes into syringe-like nano-machines that are critical for bacterial infection by many human pathogens including *Salmonella, Shigella, Yersinia* or *EPEC* (enteropathogenic *E. coli*). They actively deliver toxic bacterial effector proteins into their respective host cells ^6^. As a consequence, T3SS-mediated infection can be prevented by interfering with the functionality of the T3SS ^7^. The core of the T3SS (SPI-1, *Salmonella* pathogenicity island-1) is the needle complex (NC) ^8^, a membrane-embedded structure that mediates contact between bacterial and eukaryotic cells and contains a shielded secretion path for a safe delivery of unfolded effector proteins ^9, 10^. The needle complex is intricately assembled from multiple proteins (Supplementary Table 7) in a series of stacking membrane rings (PrgH, PrgK, InvG) to form a ‘basal body’ from which a helical needle filament (PrgI) extends into the environment ^11, 12^. The basal body encompasses the centrally located export apparatus (SpaP, SpaQ, SpaR, SpaS, InvA), which defines the entry of the secretion path into the secretion channel ^10^ and continues into the filament by attaching it with the inner rod protein PrgJ. All these proteins are thus essential for T3SS function ^7, 13^. Recent structural investigation of the evolutionary related export apparatus from the flagellar system revealed a pseudo-helical arrangement of a ternary sub-complex ^14, 15^ (FliP, FliQ, FliR (flagellar system)/SpaP, SpaQ, SpaR (injectisome), Supplementary Table 7). The FliPQR complex has been recombinantly produced in the absence of the flagellar context and the unusual packing of the individual protein chains, which all have been bioinformatically predicted as integral membrane proteins, together with the previous observation that the PQR-subcomplex precedes basal body/needle complex assembly (**‘initiation/nucleation phase’**) into functional pathogenic T3SSs (Figure 1a) ^13^ raises the question, as to how this is mechanistically controlled. Notably, ring formation of the inner (IR; PrgH, PrgK) and outer rings (OR; InvG) and their interaction into stacked rings (**‘ring-phase’**) to generate non-functional basal-body-like complexes can also occur in the absence of export apparatus proteins, suggesting that this processes is self-sufficient ^13^. Furthermore, the observation that the oligomericity of rings in the flagellar system can vary ^16^, similar to earlier reports of different ring-diameters in pathogenic T3SSs ^17^, together with increasingly high-resolution structural investigations of a defined oligomericity of the OR (15-mer) and the IR (24-mer) ^11, 12, 18^, challenges how the OR and IR specifically interact and whether and how oligomericity is controlled. Ultimately, an answer to this conundrum would clarify whether a structurally singular or diverse group of physiologically relevant T3SSs exists.

**Figure 1:**
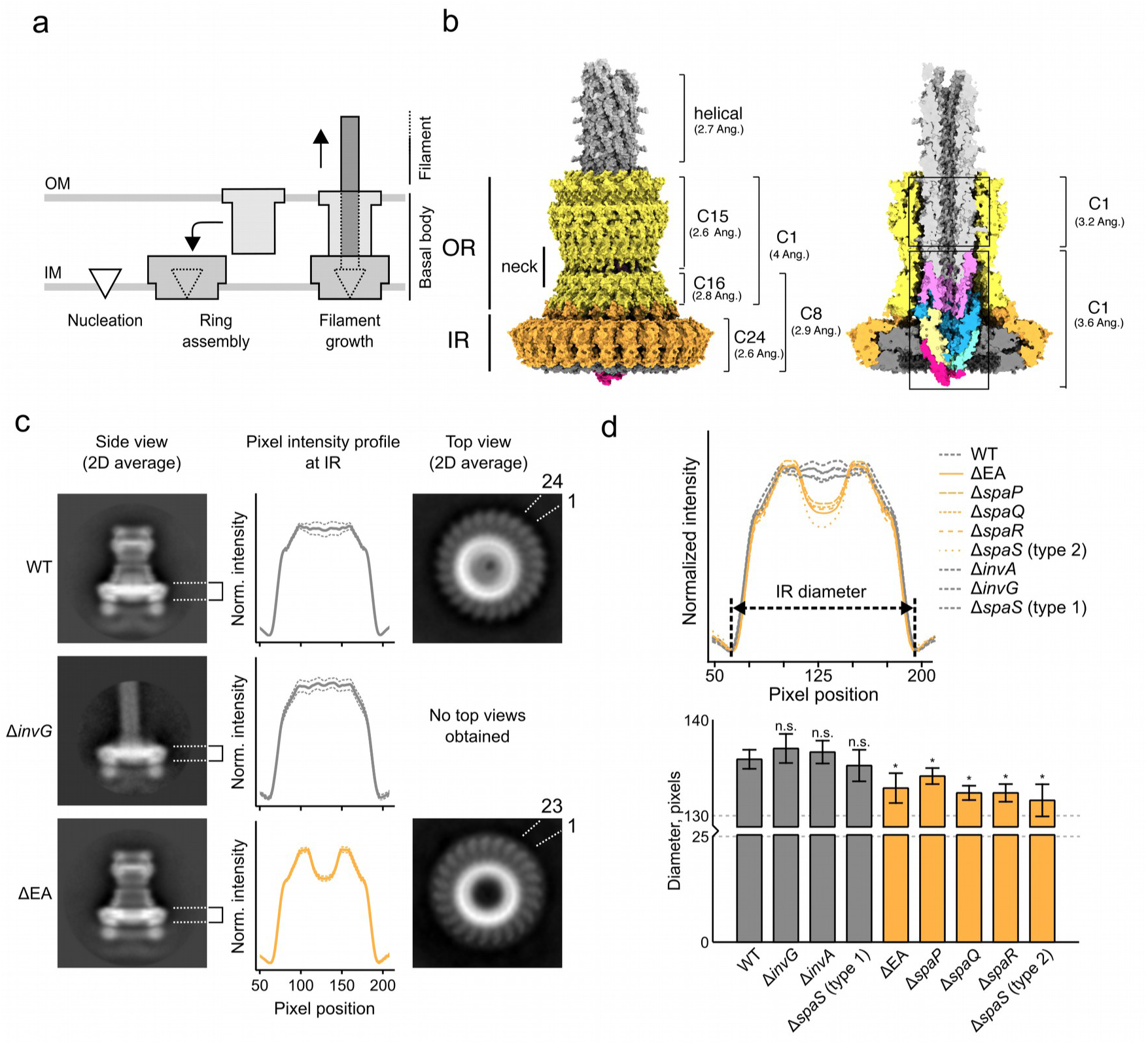
The export apparatus (EA) is essential for assembly into active complexes. a) Schematic of the overall organization and assembly phases of the T3SS needle complex. b) Overview of the needle complex architecture and identified local symmetries. c) Absence of the export apparatus components (SpaPQRS, InvA), but not OR components (InvG), results in a 23-fold symmetry of IR. Left column: 2D averages of single particle cryoEM side views of isolated WT, ΔEA needle complex but also complexes isolated from strains of individual knock-outs*invG* and ΔEA needle complex but also complexes isolated from strains of individual knock-outsEA complexes. Central column: Averaged density profile at the IR (dashed lines indicate standard deviation of averaged pixel values). Right column: 2D averages of single particle cryoEM top views of isolated WT and ΔEA needle complex but also complexes isolated from strains of individual knock-outsEA needle complexes. The number of IR protomers is indicated. d) Upper: Averaged density profile from individual knock-out needle complexes. IR diameters are measured as the distance of the minima in the density profile. Graphs of complexes showing a WT-like diameter are colored in grey. Graphs of complexes with a significantly smaller diameter are colored in orange. A reduction of normalized pixel intensity in the central area of the IR plane is visible for the same knock-outs, indicating the absence of the EA. Lower: Diameters of IR and statistical significance (*). WT-like diameters colored in grey, ΔEA-like diameters colored in orange. Complexes from a Δ*spaS* mutant could be classified into, WT-like (type1) and ΔEA-like phenotypes (type2). See also Supplementary Table 1.

Assembly of needle complexes is finalized by the growth and attachment of the filament to the export apparatus (**‘filamentous phase’**, Figure 1a). This phase is already dependent on the secretion activity of the T3SS, which transports protomers of PrgJ and PrgI into the chamber of the basal body, where they assemble into a filamentous structure that connects to the export apparatus within and extends as a PrgI-only helical filament projecting away from the basal body ^6, 17, 19^. Recent work has identified PrgJ residues that are in the vicinity of the export apparatus and a single helical turn of PrgJ has been suggested to serve as an adaptor to the filament ^20^. Nevertheless, it remains largely unknown, how monomers of PrgJ and PrgI are packaged into a quaternary structure and attached to the export apparatus, which is critical to form the majority of the protein secretion channel within the basal body.

To dissect the coordinated process of the directionality of the assembly phases into needle complexes of pathogenic T3SSs and to understand the molecular signatures promoting high-order assembly and stability, we investigated the cryo electron microscopy structures of the entire wild-type (WT) and export-apparatus (EA) deletion mutant complexes from *Salmonella enterica* serovar Typhimurium to 2.5-4 Å resolution and performed extensive mutational and functional analyses. We show that (1) oligomericity of the rings is controlled by the nucleating SpaPQR complex of the export apparatus, (2) an unusual structural pleiotropy is crucial for inter- and intra-ring stability, and (3) the synthesis of the intra-basal secretion channel is the consequence of a remarkable packing of the asymmetric export apparatus to the symmetrically helical filament. The fact that without a coordinated assembly line, highly structured basal body-like complexes can be assembled, but with a diverging quarternary architecture leading to non-functionality, emphasizes the necessity and importance of flawless assembly processes for molecular machines in general.

## Methods

All relevant materials and methods are described in the supplementary documents (genetic manipulations, secretion assays, biochemical purifications, co-variation analyses, electron microscopy and analyses, sample vitrification and cryo electron microscopy, single particle and helical reconstructions, model building, model refinement and model validation, visualization, deposition).

## Results

### The structure of the entire needle complex

We first purified WT needle complexes ^10^ from *Salmonella*, and vitrified the sample on holey carbon grids containing a layer of either continuous carbon or a freshly made graphene oxide as a sample support (Supplementary Table 3). This procedure allowed us to image particles at a wider angular distribution (Supplementary Figure 5) during cryo electron microscopy. We subsequently determined the structure of the entire needle complex by single particle analysis. Due to the size and the flexibility of the needle complex ^11^ we applied focused refinements to various parts of the complex and resolved individual sub-structures to resolutions ranging from 2.6 to 4 Å (Figure 1b). While overall the needle complex is an asymmetric molecule, individual parts of the complex display non-symmetric (C1), cyclic (C8, C15, C18) and helical symmetries.

### The export apparatus proteins dictate inner ring symmetry

Both, the IR (PrgH/PrgK) and the OR (InvG) are highly oligomeric rings that are able to self-assemble into a basal body even in the absence of the export apparatus ^13^. Because of the tight and highly stable arrangement of InvG into a 60-strand beta-barrel and its ability to independently form the outer ring in the absence of IR components ^18^, we hypothesized, whether the symmetry and the assembly of the larger IR with its 24-fold oligomeric state is governed by the InvG OR. We thus analyzed structures of purified and vitrified complexes from an InvG knock-out ^13^ and compared it to the wild type (WT) basal body complex. We found that the IR dimension from the InvG knockout is indistinguishable from the WT, suggesting that the 24-fold symmetry as clearly observed in WT from end-on views is maintained (Figure 1c). Thus, the 24-fold IR-ring formation is independent from the presence of the outer membrane protein InvG.

To determine if the centrally located export apparatus (EA) of the needle complex controls IR symmetry, we next visualized complexes isolated from an EA-knockout strain (ΔEA: Δ*spaPQRS*, Δ*invA*). Remarkably, we found that the IR in these complexes is smaller in diameter and contains only 23 subunits, as observed from end-on-views (Figure 1c). To further identify, which components of the export apparatus are responsible for this atypical structural phenotype, we analyzed previously purified and vitrified structures from individual SpaPQR, InvA knock-out strains ^21^. We found that structures obtained from an *ΔinvA* strain are WT-like, whereas structures recovered from *ΔspaP, ΔspaQ*, or *ΔspaR* strains are indistinguishable from the complete EA knock-out (Supplementary Table 1). Interestingly, the complexes obtained from the *ΔspaS* strain could be sub-classified in two phenotypes that either are WT-like (75%) or EA-knock-out-like (25%) (Figure 1d, Supplementary Figure 1, Supplementary Table 1). Together with earlier findings that PQR forms an initial complex during the assembly ^13^, our observations lead us to propose a model that the PQR sub-complex generates a structural template for the formation of precisely 24-subunits of PrgH and PrgK, respectively, for the complete IR assembly.

### The structure of the export apparatus provides a structural template for needle complex assembly into functional injectisomes

The structure of the EA bound in the center of the needle complex reveals a conical shape with a pseudo-helical arrangements of individual protein chains, displaying a P:Q:R = 5:4:1 stoichiometry (Figure 2, Supplementary Figure 8, Supplementary Table 5). The sub-structure is structurally similar to reconstituted samples of the flagellar export apparatus ^15^. The export apparatus proteins SpaP, SpaQ, SpaR, and SpaS have been bioinformatically predicted as membrane proteins containing alpha-helical hydrophobic stretches suggesting a canonical lateral organization in a lipid bilayer. However the spiral arrangement of 4 consecutive SpaQs at the bottom of the EA sub-structure generates a cork-screw-like hydrophobic belt (Figure 2c) consistent with the idea that SpaQ is in part lifted out of the membrane during the assembly^15^. A single chain of SpaR intersects the assembly of 5 successive SpaPs (Figure 2b), which conclude the upper part of the EA with a hydrophilic, acidic belt, indicating that these proteins do not behave as classical integral membrane proteins. The separation of the EA into two surrounding belts, different in their surface properties, demarcates its position relative to the surrounding PrgK ring (Figure 2c, d). At the corresponding position of the PrgK ring are two complementary enclosures established by a lower and conserved, largely hydrophobic (aspartate-92 to proline-98, Figure 2d,e (arrow)) and a higher hydrophilic loop (glycine-137 to proline-142) that project towards the EA. Starting from the bottom tip of the PQR conus, the PQR complex is slightly located off center relative to the projected center determined by the PrgK ring. From there, it spirals upwards and establishes hydrophobic contacts primarily to the lower PrgK-loop (Figure 2d, lower, Supplementary Figure 2). Subsequent interactions to PrgK are determined by the curvature of the PQR-complex and are captured largely within the first half-circle of PrgK, explaining how the PQR complex provides a structural template for ring assembly. The interaction to PrgK is further stabilized by contacts of SpaP, which constitutes the upper part of the EA. Interestingly, the fourth component, SpaS, is not visible in the reconstructions suggesting that it has been largely stripped off during the purification. This is in agreement with earlier biochemical data ^13^. Nevertheless, because SpaS only fractionally influences the efficiency of proper ring assembly, we considered that SpaS is localized at the PQR complex in the vicinity of the PrgK ring position. We thus performed a co-variation analysis, showing that indeed the strongest signals are obtained between SpaS-SpaQ and SpaS-SpaR (Supplementary Table 2). We used this information together with a template structure from a flagellar EA complex (unpublished data, S. Lea) to model SpaS onto the PQR complex (Figure 2f), demonstrating that the majority of SpaS is positioned along a conserved patch of the hydrophobic belt (Figure 2g) and surrounds the tip of the hydrophilic PQR conus. However, two SpaS loops reach into the conserved hydrophobic PrgK enclosure, thereby completing a circular packing of PQRS in the center of the PrgK ring (Figure 2d).

**Figure 2:**
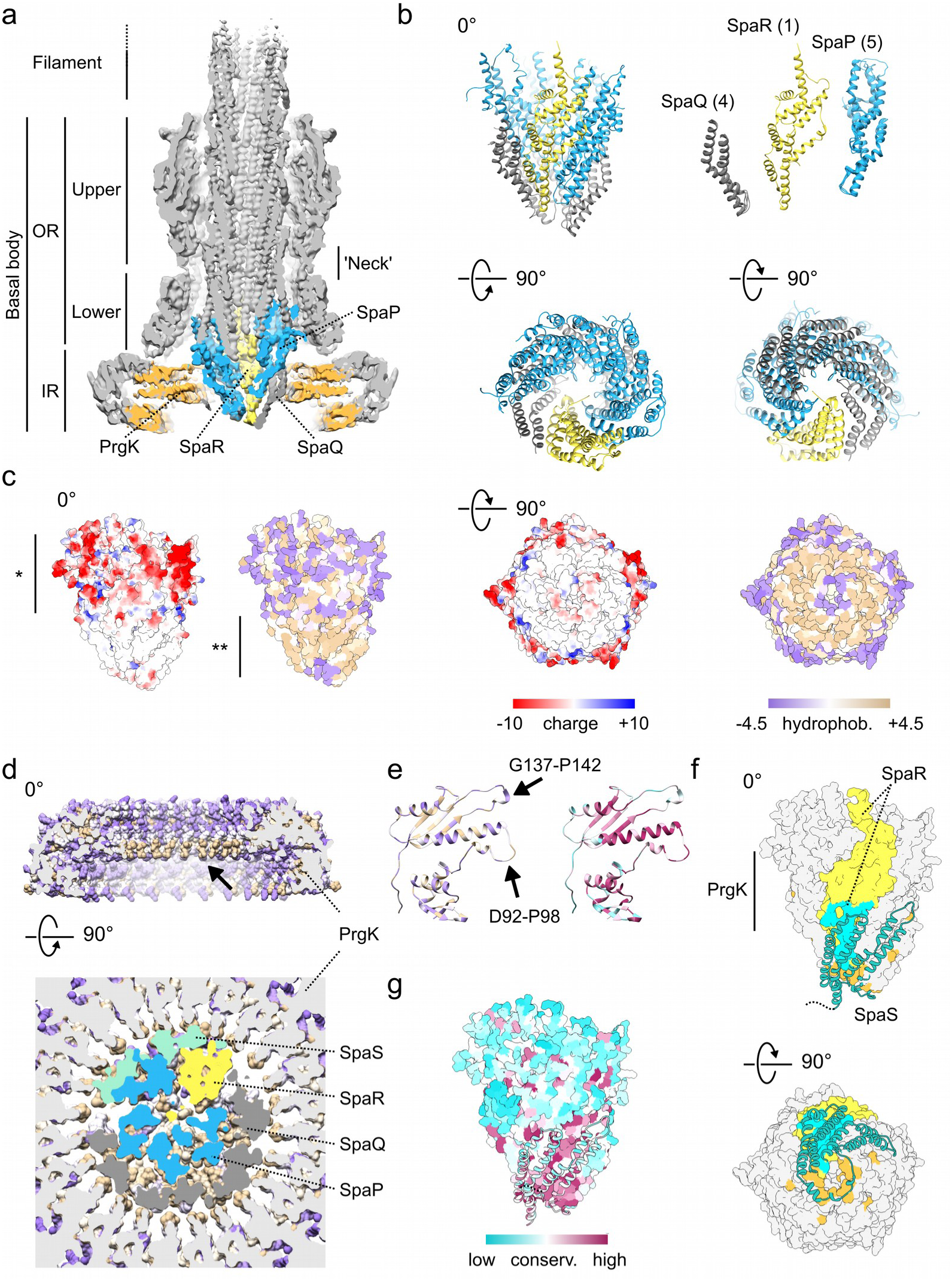
The structure of export apparatus within the needle complex reveals a structural template for ring formation of active complexes. a) Central cross-section of asymmetric (C1) single-particle reconstruction data of WT needle complexes. OR: outer ring, IR: inner ring. Export apparatus components are colored in dark grey (SpaQ), yellow (SpaR) and dark cyan (SpaP), respectively. Densities for PrgK are colored in orange. b) Structure of the PQR complex present in needle complexes shown as a ribbon diagram from different views (side, top, bottom). The PQR complex adopts a pseudo-helical arrangement with the sequential order of 4x SpaQ (dark grey), SpaR (yellow), and 5x SpaP. (Top right) Individual protein chains for SpaQ and SpaP, respectively, are structurally similar to each other (top-right corner). RMSD range for SpaQ: 1.3-2.4 Å, RMSD range for SpaP: 1.5-3.8 Å (see Supplementary Table 5). c) The surface of the PQR complex contains a charged (*) and a hydrophobic (**) belt. The electrostatic potential was calculated according Coulomb’s law (dielectric constant ε=4). The hydrophobicity is colored according to the scale of Kyte and Doolittle ^30^. Side views on the left are rotated around the x-axis to provide bottom-views of the EA on the right. d) Upper: half-cut side view of the PrgK ring shows a hydrophobic and conserved loop facing the interior of IR (arrow). Lower: The hydrophobic belt of the PQR complex binds tightly to the surrounding hydrophobic PrgK ring. e) The loop (aspartate-92 to proline-98) of PrgK is largely hydrophobic and conserved. f) Homology model of SpaS (ribbon) bound to the PQR complex (surface). Evolutionary couplings between SpaS/SpaR and SpaS/SpaQ are shown in cyan and orange, respectively. Notably, SpaS twists in a circle around the conical PQR entry. g) Surface representation of the PQR(S) complex colored by conservation as determined by ConSurf ^31^.

Taken together, the structure of the EA explains how the PQR/S-sub-structure can serve as a structural template for inner ring formation into physiologically relevant 24-mer complexes. This is also consistent with earlier biochemical observations that the export apparatus, in particular PQR, followed by PQRS forms a sub-structure that initiates assembly into full WT basal body^13^. Our structure of the EA within fully assembled injectisomes also clarifies that the majority of PQR (and S) monomers are embedded outside a membrane bilayer and thus represent a class of molecules that either can be membrane proteins during assembly or soluble proteins within the complex.

### Active injectisomes are asymmetric and structurally pleiotropic

Entire needle complexes are structurally complex due to their size, their stoichiometry and copy-number of polypeptide chains, their various symmetries (cyclic, helical) and asymmetry (pseudohelical conus) in sub-parts of the injectisome. As a consequence, many interactions at and between various parts are necessary to establish sufficient stability but also conformational flexibility, essential for assembly ^17^. To address the relevant factors for proper needle complex assembly, we first analyzed the interaction of the larger inner (PrgH/K) and smaller outer membrane ring (InvG) by focused reconstructions at the interface of the IR and OR (Figure 3, Supplementary Figures 9-10). We found that 16 out of 24 PrgH monomers tightly associate with their ∼30 amino acid C-terminal tails to 16 subunits of the N-terminal InvG-ring (domains N0-N1). They form 8×2 beta-sheets (conformation “B’ and ‘C’ in PrgH) that intercalate between 16 beta-stranded InvG domains and thus establish together with InvG a large circular and flat beta-stranded ring facing the 24-mer PrgH/K ring (Figure 3a). Intercalation and formation of the C-terminal beta-sheet is essential to confer stability between the rings (Figure 3b) and explains previous functional data of truncated PrgH ^22, 23^. Furthermore, the remaining 8 subunits of PrgH also form a short two-stranded beta-sheet (conformation ‘A’), which binds to the exterior part of InvG as well as to a neighboring PrgH and presumably stabilizes the overall arrangement.

**Figure 3:**
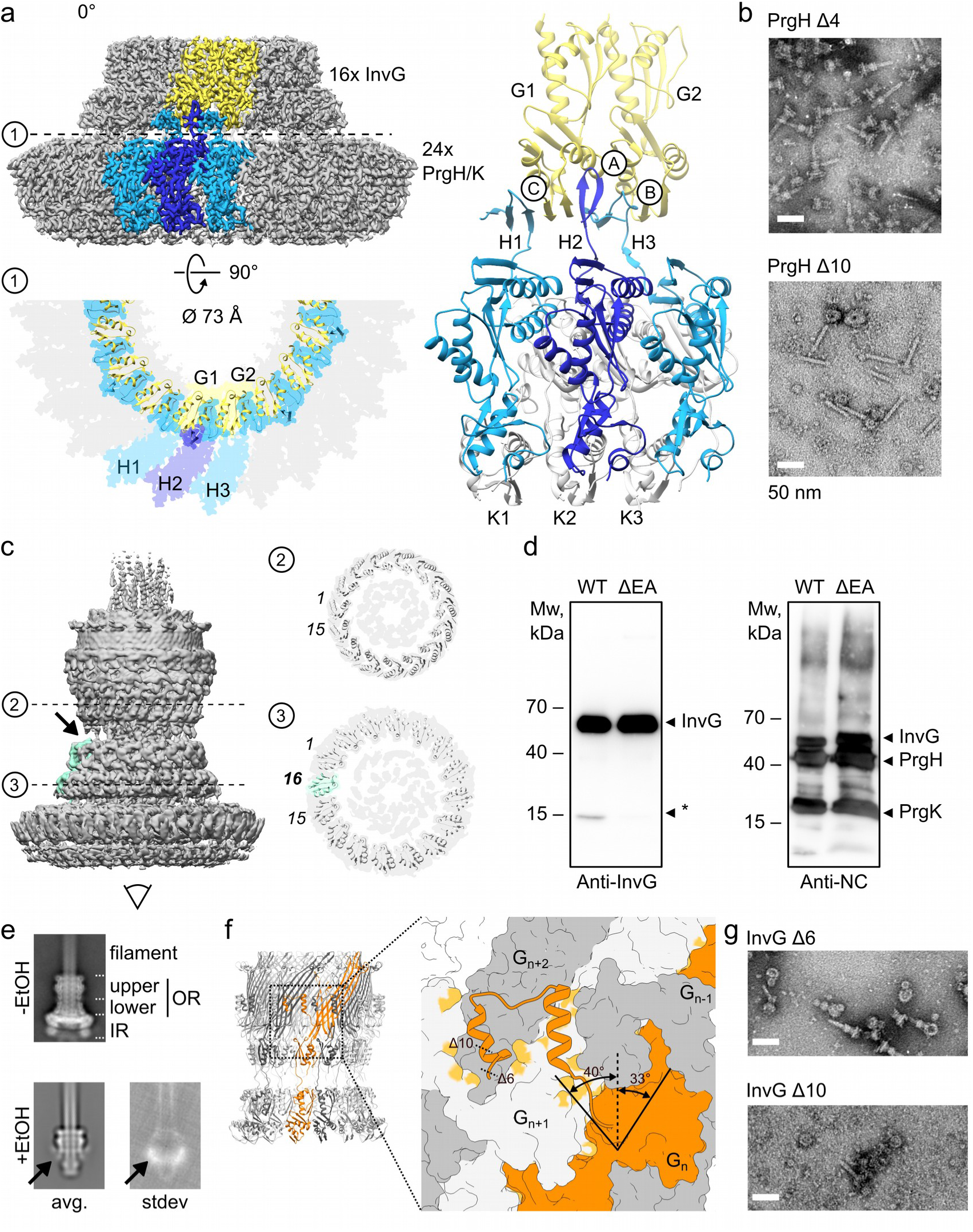
WT needle complexes are asymmetric and structurally pleiotropic. a) Cryo electron microscopy structure of the IR and OR of WT needle complexes. Twenty four protomers of PrgH and PrgK, respectively, interact with sixteen InvG protomers. The interaction is established by three repeating conformations at the C-terminal tail (amino acids 361-392, conformation ‘A’, ‘B, ‘C’) of PrgH resulting in an extended circular beta-sheet composed of PrgH and InvG. The overall structure displays a C8 symmetry with 3 PrgH (H1, H2, H3, blue), 3 PrgK (K1, K2, K3, grey), and 2 InvG (G1, G2, yellow) in the asymmetric unit. b) Successive truncations of amino acids at the PrgH C-terminus impair complex formation. Deletions of ten amino acids do not yield stable needle complexes as analyzed by electron microscopy. Scale bar 50 nm. See also Supplementary Figures 3 and 6. c) The OR in WT needle complexes assembles into a lower 16-mer, and an upper 15-mer ring complex (C16/C15) that are slightly tilted towards each other. The 16^th^ protomer of InvG in the lower ring is highlighted in green (C1 reconstruction). The connecting densities at the furthest distance at the constriction are missing on one side (arrow). Cross section of the upper and lower OR at the indicated positions. d) Western blot detection of a faster migrating InvG band in isolated WT needle complexes using a polyclonal antibody raised against the N0-N1 domains of InvG. The band is not detected in ΔEA mutant complexes. Other needle complex components are detected using an anti needle complex antibody. e) Mild and selective needle complex disassembly. Class averages of vitrified samples prior and after ethanol treatment. In the absence of the IR, the lower OR does not maintain a ring conformation, whereas the upper OR stays intact. (avg (average) and stdev (standard deviation of class average). See also Supplementary Figures 3 and 6. f) g) The C-terminus of InvG spans two neighboring InvG subunits and is critical for injectisome stability. Ribbon diagram of the entire 16/15-mer OR and zoomed in view in surface view. A single InvG (orange) interacts with two consecutive InvG protomers (G_n+1_, G_n+2_ light grey and dark grey, respectively). Positions of truncations (Δ6 and Δ10) impacting complex assembly (g) are indicated with dashed line. Scale bar 50 nm. See also Supplementary Figures 3 and 6.

The 16-mer structure of the secretin InvG is in sharp contrast to previous high resolution structures that reported 15 subunits for this domain ^12^. Interestingly, only the upper, C-terminal domain of InvG maintains a 15-mer ring, as shown in our 2.6 Å resolution structure (Supplementary Figure 11, Table 4) and similar to earlier reports ^18^. This raises the question as to how the individual InvG monomers that start with their N-terminal domain linked to the larger inner ring span the periplasm and extend into a the 15-mer upper OM ring. We thus performed focused reconstructions in C1 of the full OR (Figure 3c). At first, we found that the upper and lower domains of the OR are slightly tilted and that the continuous density bridging the constricted ‘neck-region’ (labelled in Figure 2a) is largely present only on one side of the neck. Notably, we found that indeed, the entire OR assembles into a lower 16-mer and an upper 15-mer ring, suggesting that the upper part of the16th subunit is left out during the 15-mer ring assembly. Moreover, we found that the 16th InvG monomer is cleaved in isolated needle complexes, as demonstrated by the detection of a faster migrating band in Western blots using an antibody raised exclusively against the lower InvG-ring domains (N0-N1, Figure 3d). To test, whether the lower OR, represented by the N0-N1 domains of InvG, remains stable as a ring in the absence of the larger IR, as an indicator for stacking of preformed rings^24^, we established a mild disassembly protocol that selectively removes the IR components. Surprisingly, we found by cryo electron microscopy that the lower OR (N0-N1) becomes largely unstructured and the individual domains very flexible, whereas the upper secretin ring maintains as a stable ring (Figure 3e). Similar observations have been reported upon isolation of only secretin-rings ^18^. This demonstrates that the so-called ‘ring-forming’ domains N0-N1 ^12^ in the lower part of InvG are in fact unable to independently form stable rings and require the presence of the larger 24-mer inner membrane ring for (16-mer) ring formation. Consequently, we speculated, whether the stability of the entire secretin ring is largely mediated by the overall structural arrangement of the upper InvG ring. While the majority of the individual monomers are arranged side-by-side, generating a beta barrel with 60 strands tilted by +33°, a long C-terminal stretch (amino acids: V519-G557) interacts as an extended conformation at approximately −40° with two neighboring InvG monomers in the opposite direction (n-1, n-2) of the tilted strands (Figure 3f). To test, whether the C-terminal end of InvG indeed provides stability for upper ring formation, we generated C-terminal truncation mutants, probed for functionality and analyzed complex formation (Figure 3g). We found that truncations of only six amino acids already impacted, and ten amino acids that eliminate only a fraction of the very last alpha helix of InvG, completely abrogated needle complex formation and function (Figure 3g, Supplementary Figures 3 and 6).

Taken together, the InvG ring in functional needle complexes is structurally pleiotropic, displaying different stoichiometries in the upper and lower OR. The assembly of the secretin into functional needle complexes requires 16 monomers at the basal side to interact with 8×2 intercalating PrgH C-terminal domains (CTD) at the inner membrane. It is furthermore stabilized by binding of 8 additional ‘exterior’ PrgH-CTD’s, and 15 InvG protomers to form a stable ring at the apical side that connects to the outer membrane.

### The atomic structure of a dead-end T3SS

The fact that complexes obtained from EA knock-outs exclusively adopt a 23-mer arrangement in the larger IR (Figure 1c) from otherwise native PrgH/PrgK proteins raises the question, as to how basal-body formation is structurally controlled. In order to gain detailed structural insight into the mechanism by which assembly into aberrant injectisomes diverges from assembly into functional systems, we determined the atomic structure of the export apparatus knock-out T3SS (Figure 4). We thus applied a similar purification strategy and observed by negative-stain electron microscopy that the entire ΔEA needle complex but also complexes isolated from strains of individual knock-outsEA needle complex but also complexes isolated from strains of individual knock-outs for SpaP, SpaQ, and SpaR are less stable (Supplementary Figure 7). Very often, the larger inner and smaller outer membrane rings were displaced from the complexes.

**Figure 4:**
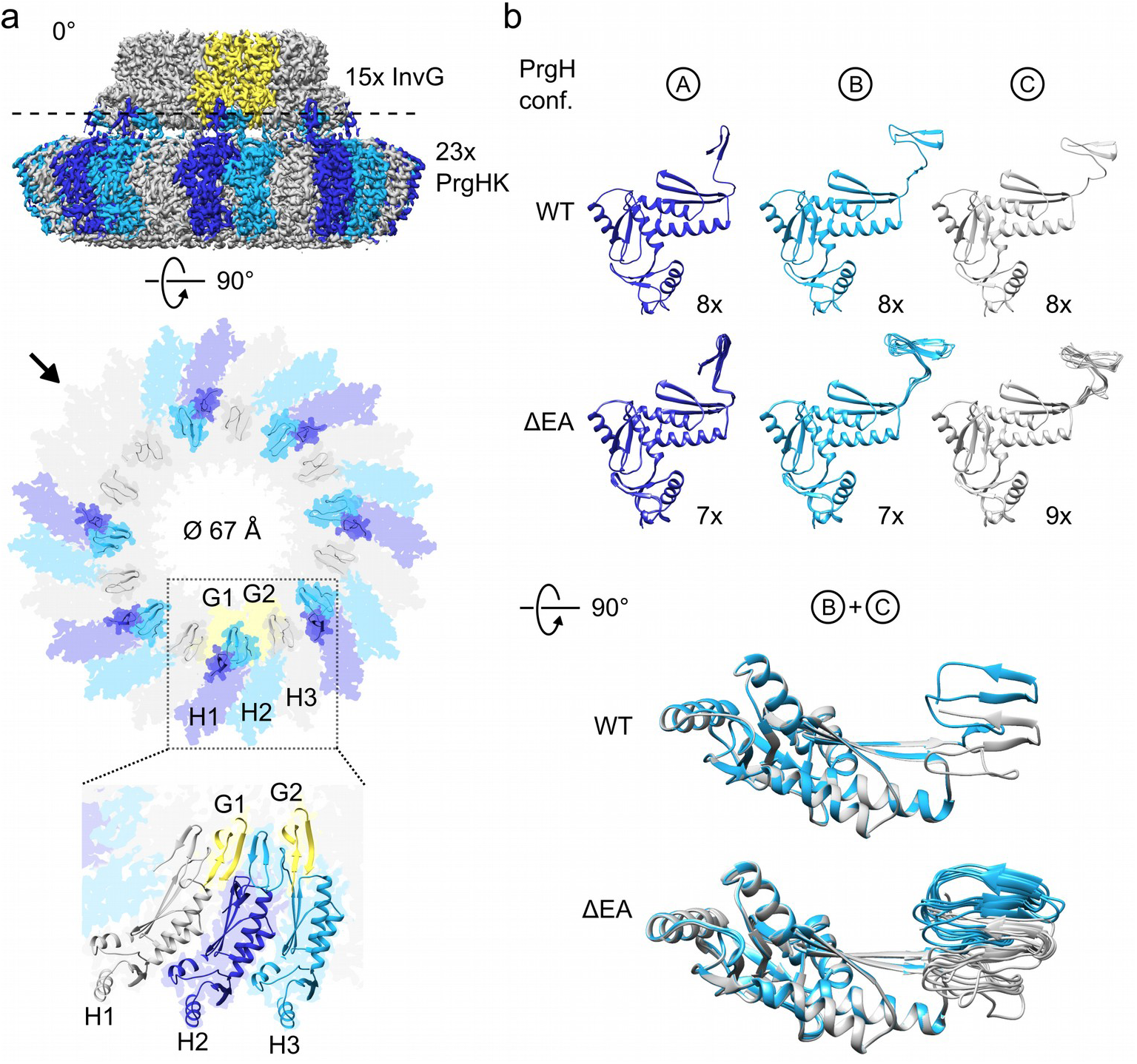
The structure of dead-end complexes. a) Cryo electron microscopy structure of complexes isolated from a ΔEA needle complex but also complexes isolated from strains of individual knock-outsEA strain. focused reconstruction of the IR/OR (lower) in C1 revealed a 23:15 symmetry of IR:OR. Lower: ribbon diagram of the IR/OR interaction. PrgH protomers showing the same conformation at the C-terminal tails are indicated with the same colors. Zoomed view: three protomers of PrgH with different conformations (H1, H2, H3), two protomers of InvG (G1, G2). The symmetry break occurs at the position indicated with an arrow. b) Three conformations of PrgH (‘A’, ‘B’, ‘C’) differ at the C-terminal tail of PrgH in WT and in ΔEA needle complex but also complexes isolated from strains of individual knock-outsEA complexes, respectively. Conformation ‘A’ binds to the outside of InvG, whereas ‘B’ and ‘C’ intercalates between the N-terminal domains of InvG. In WT the 3 conformations are arranged symmetrically (24 protomers: 8x each conformation), whereas in ΔEA needle complex but also complexes isolated from strains of individual knock-outsEA complexes 7x conformation ’A’, 7x ‘B’, and 9x ‘C’ are arranged within the 23-mer PrgH ring. Overall, the intercalating conformations ‘B’ and ‘C’ (362-392) have the same structural fold but differ in their lateral position relative to the main structure of PrgH.

Interestingly, this behavior was largely absent upon vitrification, indicating that negative staining induces sufficient shear forces that induce structural deviations in isolated knock-out but not WT complexes. We solved the cryo electron microscopy structure from the ΔEA needle complex but also complexes isolated from strains of individual knock-outsEA basal-body complex to 2.5-3.4 Å (C23; C1) resolution (Table 4). Overall, the PrgH/ K ring adopts a 23-mer structure with a similar intertwined arrangement of non-tilted PrgH and 24° tilted PrgK (Supplementary Figure 4a) that stabilizes the large IR. Comparison of the interacting surfaces of neighboring (PrgH,PrgK):(PrgH,PrgK) protomers revealed a larger area in the ΔEA needle complex but also complexes isolated from strains of individual knock-outsEA 23-mer than in WT (23-mer: 4526.5 Å^2^; 24-mer (WT): 4435.5 Å^2^).

Overall, the 23-mer ring has a smaller diameter (inner diameter 67Å) than in the WT complex (inner diameter 73Å), with insufficient space for the EA located at the equivalent position and height relative to the PrgK rings (Figure 4, Supplementary Figure 4b). This together with the previous packing of EA in WT complexes, establishes that the EA plays an essential role during assembly by generating a structural template that mechanistically counteracts the self-assembly propensity into rings and thus ensures formation of 24-mer rings.

To investigate the structural consequence of ring stacking to a 23-mer IR in respect to the interaction with the OM ring, we determined the upper and lower InvG ring from the ΔEA needle complex but also complexes isolated from strains of individual knock-outsEA complex by focused reconstructions to a resolution of 2.7 and 2.8 Å, respectively. To our surprise, we found only a 15/15-mer arrangement of the secretin (Supplementary Figures 17 and 18). This is in sharp contrast to the 16/15-mer arrangement in WT complexes (Figure 3c). This observation is also consistent with the lack of detection of a faster migrating InvG band in Western blots (Figure 3d), suggesting that overall, the secretin protein InvG is able to adopt different configurations (15/15 and 16/15) dependent on the oligomericity of the larger inner ring. In pursuit of understanding the architecture of how a 23-mer ring interacts with a 15-mer to still being able to form mutant basal bodies, we solved the structure without symmetry enforcement (C1) (Figure 4a, Supplementary Figure 16). Contrary to WT complexes that show a highly organized and repeated pattern at the IR/OR interface (8x conformation of ‘A’, ‘B’, ‘C’, respectively, see Figure 4b), the interface of the ΔEA needle complex but also complexes isolated from strains of individual knock-outsEA needle complex is irregular and asymmetric throughout the interaction circle (conformation ‘A’ (7x), ‘B’ (7x), ‘C’ (9x), Figure 4b). Despite the suboptimal structural arrangement of the interaction surface, it is still surprising that basal body-like complexes can be formed suggesting that avidity also plays a role between ring-stabilization.

### Structure of the inner rod and internal filament

The final step for needle complex assembly requires active transport of the inner rod protein PrgJ and the needle filament PrgI, which generate a filamentous sub-structure for the secretion of later substrates by connecting the pseudo-helical packing of the conical EA to the exterior helical needle filament structure (Figure 5a). Both proteins are small (101 residues in PrgJ, 80 residues in PrgI) and predominantly alpha helical in nature. To clarify, how PrgJ and PrgI connect and are arranged within needle complexes, we solved the cryo electron microscopy structure by focused refinements around the central part of the needle complex (Figure 3b, Table 4). We found that PrgJ adopts a two-legged alpha-helical structure with similar lengths (Figure 5b). Six PrgJ subunits are circularly, yet asymmetrically organized (for a single PrgJ only one alpha helix (*) is visible; blue ribbon) and interact with the export apparatus protein SpaP. The arrangement establishes a space that is subsequently occupied by PrgI (yellow), allowing individual monomers to enter between and grow upwards as a helix. The helical parameters of the internal part of PrgI are similar to the reconstructions obtained from the external filament (twist/rise 63.3°/4.4 Å in outer filament, 63.4°/4.3 Å in inner filament, Supplementary Figures 14 and 15). PrgI also establishes contacts to the surrounding InvG secretin ring (Figure 5b), suggesting a firm stabilization of the filament within the basal body. Notably, the first five PrgI monomers (yellow) that are localized between the PrgJ monomers are partly unstructured at amino acids 1-20 in contrast to later PrgI subunits throughout the helical assembly, suggesting that this is a crucial structure that allows the transition from a non-helical to a helical arrangement to occur.

**Figure 5:**
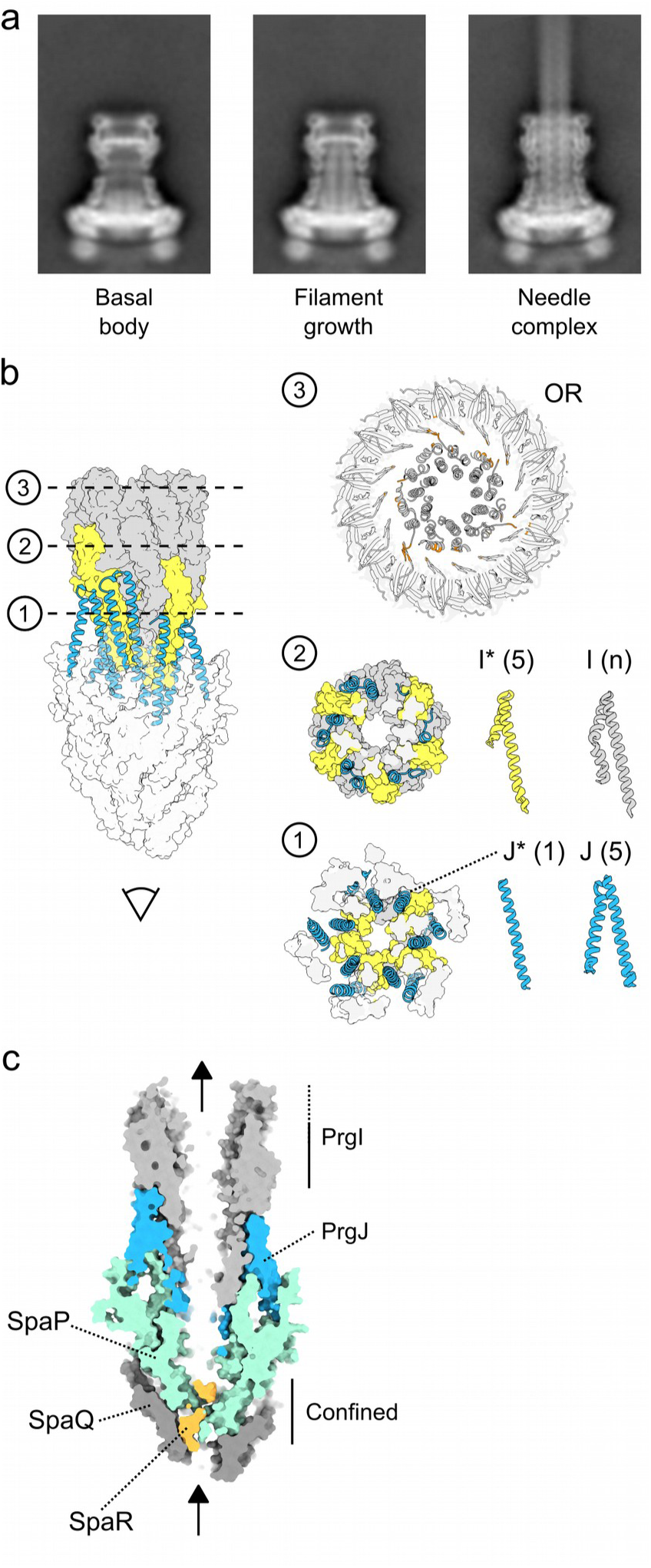
The PQR complex, PrgJ and PrgI form an intricate secretion path for the substrate. a) Distinct class averages of vitrified samples isolated from a WT resemble the transition from a basal body to the fully assembled needle complex. The internally growing filament pushes towards the septum. In assembled needle complexes, the septum underwent a conformational change to allow extra-basal polymerisation of the needle filament. b) Structure of the entire internal sub-structure. Six PrgJ and the initial five PrgI protomers serve as adaptors from pseudo-helical to a true-helical symmetry of the needle filament. Left: Overview of the EA to needle filament structure, shown as a surface side view with EA components in white, PrgJ subunits in blue ribbons representation and interlocked PrgIs in yellow (surface). Three cross-sections show the packing of PrgJ connecting to the EA (‘1’), PrgI adaptors (yellow), and the helical PrgI filament stabilized by interactions (orange) to the surrounding upper InvG ring. (InvG ring not shown in side view). PrgJ forms two-legged helix-turn-helix structures that serve as pillars to support the filament (gray). (2) The helical arrangement of PrgJ is adapted by a single round of five PrgI protomers (yellow), with a distorted N-terminal region (I*) similar to the distortion observed at one N-terminal helical leg of a single PrgJ (J*) protomer. (3) The now purely symmetrical filament extends upwards within the OR InvG N-terminal secretin domain. Interacting residues of PrgI (N-termini) and InvG protomers are colored in orange. All proteins are shown in cartoon representation. c) Substrate path through the EA, PrgJ, and the PrgI filament. The EA confines the substrate path that continues at the level of SpaP into a 12-16 Å wide channel.

Secretion of unfolded substrates occurs through the export apparatus and continues into the filamentous sub-structure. We thus analyzed the entire channel and found that it starts with a hydrophilic ring at its conical entry (Figure 5c, arrow). Shortly after, the channel snakes through a highly confined area, to then reach a space of approximately 12 Å in diameter, defined by the upper part of the export apparatus. Then, the channel continues through the filament with a slightly larger diameter (14-16 Å), similar to the dimensions observed in the external helical filament (16 Å). The highly confined areas are consistent with a mechanism that restricts passage of only completely unfolded proteins shortly after entry into the export apparatus ^10^.

## Discussion

Correct and spatio-temporal assembly of molecular machines is a prerequisite for their functioning within the cell. Here we have analyzed the molecular signatures required for the directional assembly of the megadalton-sized needle complex of pathogenic T3SSs and have elucidated requirements to establish stability among the proteins involved. We solved the structure of the prominent needle complex of the T3SS from *Salmonella enterica* serovar Typhimurium composed of > 200 polypeptides by cryoEM including co-variational data. Our work provides the molecular basis of the requirement to form a functional complex and establishes a framework for a detailed analysis of the highly coordinated and structurally controlled assembly path. Comparison with dead-end complexes including mutational and functional data allows us to propose a model for the three consecutive assembly steps, starting from an initiating nucleation phase, progression through the ring-phase, to finalize the process in the filamentous phase.

We found that a tripartite PQR complex of the export apparatus located centrally within injectisomes generates a structure very similar to isolated and recombinantly produced flagellar export complexes ^14^. This suggests that the PQR complex can form independently during the initiation/nucleation phase prior to ring-formation (Figure 6), which is also consistent with earlier biochemical findings ^13^. A defined stoichiometry of PQR (5:4:1) is essential, as the resulting pseudo-helical, conical structure resembles a template for the lateral packing of exactly 24 monomers of PrgH and PrgK, the components of the larger inner ring. The export apparatus protein InvA is not involved in correct 24-mer ring formation, consistent with its physical location away from the core of the PQR complex revealed by cryo electron tomography ^12^. However, SpaS increases the efficiency of precise assembly likely through its direct interaction at PQR and its immediate vicinity to PrgK. Indeed, the observation that in the absence of the PQR complex, the IR self-assembles to a 23-mer ring only and subsequently culminates in non-functional complexes, highlights the importance for a precise, structure-based control mechanism.

**Figure 6:**
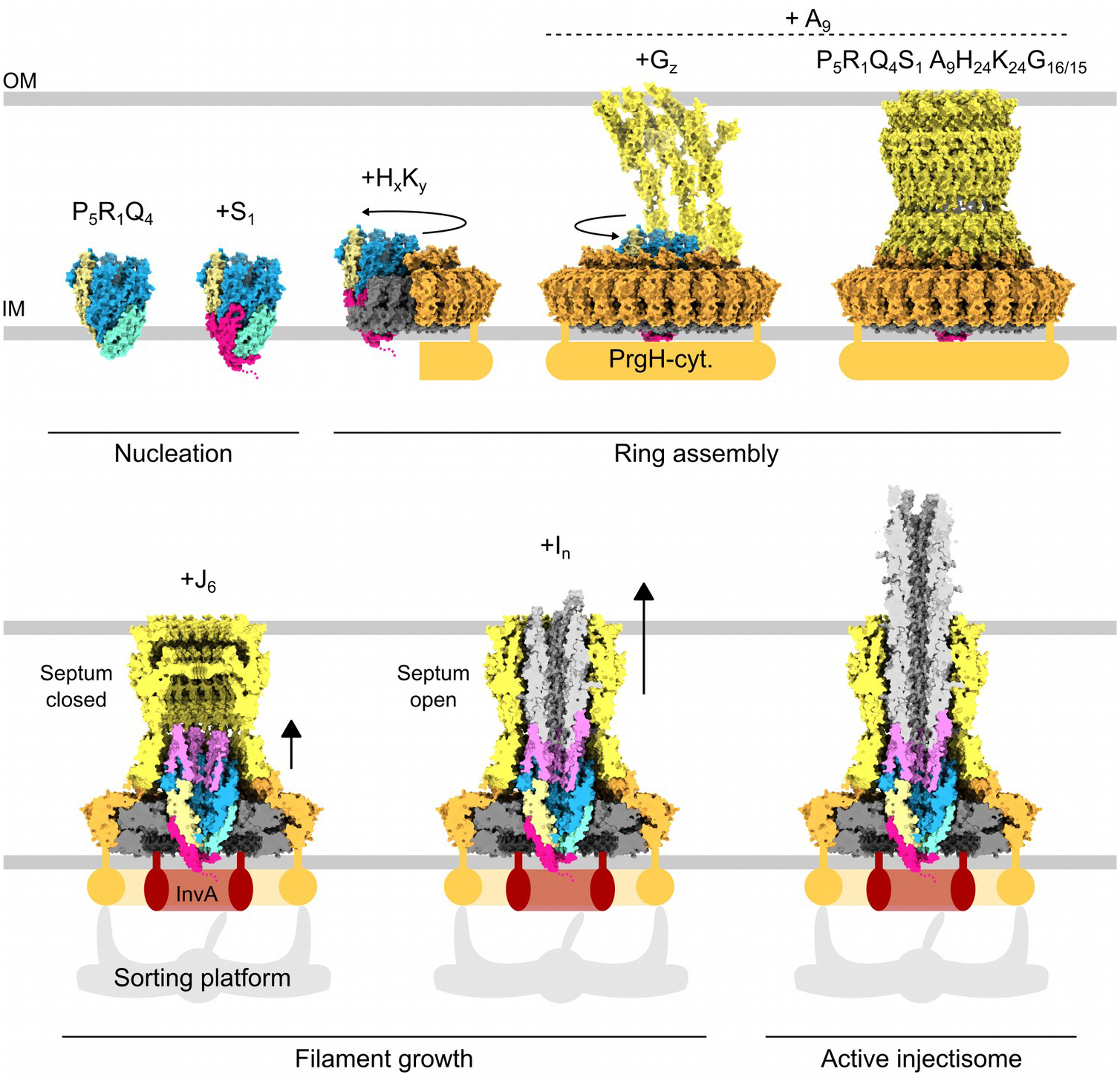
Model of step-wise assembly of injectisomes. Assembly of injectisomes is initiated in the nucleation phase that requires the export apparatus to form the PQRS complex. During ring assembly, first the inner ring components PrgH and PrgK assemble to provide an interaction point for the outer ring protein InvG. Sixteen InvG molecules bind with their N-terminal domains to 24 PrgH tails but only fifteen C-terminal domain of InvG protomers arrange into a stable outer ring excluding the 16^th^ InvG monomer. Subsequently, the filamentous phase is characterized by the growth of the internal secretion channel. PrgI and PrgJ are actively transported by the T3SS and subsequently grow a filamentous sub-structure attached to the export apparatus. Transport of PrgJ and PrgI is dependent on the recruitment of the transiently interacting sorting platform harboring, among other components the T3SS-specific ATPase. During the growth of the inner PrgI filament, the closed septum in basal bodies undergoes a conformational change to allow extra-basal growth of the needle filament to continue. (IM – inner membrane; OR – outer membrane). SpaP – blue, SpaR – yellow, SpaQ – turquoise, SpaS – magenta, PrgK – grey, PrgH – orange, InvG – yellow, PrgJ – pink, PrgI – light grey)

Similarly, the high propensity for self-assembly of InvG protomers into the upper part of the OR, mediated by a highly intertwined arrangement of individual InvG monomers, and the inability of the N0-N1 InvG domain to maintain a stable ring structure for subsequent stacking to the larger IR to establish the basal body ^24, 25^, cannot be reconciled with the here found and unusual pleiotropic structure of the OR (16/15-mer) in isolated injectisomes. Contrary to a stacking model of pre-formed rings, our data convey that 16 individual InvG protomers first connect with their N-terminal domains to the IR, allowing the distant C-terminal domain of InvG to stabilize into a 15-mer with a tightly interwoven OR 60-stranded beta-barrel. It remains unclear, why the 16^th^ InvG of the OR is excluded from the upper part of the ring, resulting in a 16/15-mer configuration within functional injectisomes. It could, however, be explained by an evolutionary re-adaptation of existing secretin folds for stable ring formation employed in other systems, such as various 15/15-mer type-2 secretion systems ^26–28^, yet constrained by the presence of a 24-mer IR. This hypothesis is also in agreement with the fact that InvG is able to adapt a 15/15-mer configuration in isolated ΔEA needle complex but also complexes isolated from strains of individual knock-outsEA complexes. Taken together, this corroborates the idea that a coordinated, structurally constrained assembly process contains the self-assembly propensity of the outer ring component.

Finally, the growth of the filamentous sub-structure requires to connect to the export apparatus and to transition from a pseudohelical export apparatus rim to the actual helical needle filament. This transition is accomplished by a highly conserved, elegant packing of the inner rod protein PrgJ to SpaP on one side and to the needle filament proteins, which are packaged in between individual PrgJ protomers on the other side.

Polymerization of PrgI protomers leads to elongation of the needle filament that pushes towards the septum of the OR. Subsequently, a conformational change is required to induce the opening of the septum and to allow the continuation of the needle filament growth into the extracellular space ^12^. There, it serves as the connecting tube that allows the safe translocation of bacterial effector proteins into host cells. The observation that the PrgI filament structure within and outside of the basal body slightly differs, supports the idea of signal transduction mediated by a conformational change through the filament upon host cell contact ^29^. Therefore, the structure of the entire needle complex presented in this study provides the basis for analyzing the mechanism of activation for secretion of toxic bacterial effectors in the future.

## Acknowledgments

This project was supported by funds available to TCM at the Institute of Structural and Systems Biology at the University Medical Center Hamburg-Eppendorf, the Institute of Molecular Biotechnology (IMBA) of the Austrian Academy of Sciences, and the Research Institue of Molecular Pathology (IMP). TCM (and SM) received funding through grant I 2408-B22 furnished by the Austrian Science Fund (FWF).

VK was supported by Boehringer Ingelheim Fonds PhD fellowship. High-performance computing was possible through access to the HPC at DESY/Hamburg (Germany). and the Vienna Scientific Cluster (Austria). Electron microscopy was preformed at the CSSB Cryo-EM Multiuser facility (Krios, L120C) and the Vienna Biocenter Core Facilites (Polara, Morgagni).

## Author contributions

NGM, VK, MJB, JM, JW, SM, OV, WL, SW, FD, TCM - designed experiments

NGM, JM, SW - prepared knockouts and mutants

NGM, VK, MJB, OV, SM, JM - purified injectisomes

NGM, JM, OV, SM, VK - biochemical tests

MJB, JW, WL - cryoEM data collection and image data management

MJB, VK, TCM - single particle and helical image data processing

VK - Co-variation analysis

NGM - de-novo building and validation

MJB, NGM, VK, LK, WL, FD, TCM - model building, refinements and interpretation

VK - prepared figures

SL, TCM - supervised model refinements

TCM – original draft

all authors – draft review and editing

TCM - supervised project

## Supplementary Methods

### Methods

#### Generation of knockout strains

The InvG knockout was created following the protocol of ^1^. The overlapping sites on the knockout cassette were designed to substitute base pairs 8 to 1446 of the *invG* gene (*Salmonella enterica* serovar Typhimurium str. LT2, AE006468.2, bps 3,041,604 to 3,043,292) to chloramphenicol resistance gene. The last 246 bps of *invG* were left intact to preserve the ribosomal binding site (RBS) for the next gene in the *inv* operon, thus avoiding polar effects of the deletion. The modified genomic region was subsequently transduced to a clean background strain (SB905) ^2^ by the *Salmonella*-specific phage P22. To remove the antibiotic resistance, the strain was transformed with the thermosensitive plasmid pCP20, containing the yeast *flp* recombinase gene, which removed the antibiotic resistance cassette via flanking FRT sites. The strain was cured from the plasmid by overnight incubation at 42°C. Knock-out strains for the entire export apparatus (Δ*spaPQRS*, Δ*invA)* and the individual export apparatus genes (Δ*spaP,* Δ*spaQ,* Δ*spaR*, Δ*spaS,* Δ*invA)* have been described previously ^3^.

Bacterial strains and phages used in this study are summarized in Supplementary Table 6.

#### Complementation plasmids

The *ΔinvG Salmonella invG Salmonella* strain (SB908) was complemented with WHS008 generated by replacing the PrgH sequence in the plasmid WHS006 ^4^. Additionally, a stop codon was introduced to avoid expression of the polyhistidine-tag.

C-terminal deletion variants of the complementation plasmids WHS006 (for Δ*prgH* complementation) and WHS008 (for *ΔinvG Salmonella invG* complementation) were generated by an inverse site-directed mutagenesis protocol utilizing and checked by Sanger sequencing.

The plasmids were electroporated into *ΔinvG Salmonella prgH* or *ΔinvG Salmonella invG Salmonella* strains carrying the additional plasmid pSB1418, carrying *hil*A gene, the main transcriptional regulator of SPI-1 ^5^ controlled by the arabinose promoter.

Plasmids used in this study are summarized in Supplementary Table 6.

#### Secretion assay

Secretion assays were conducted as previously described ^6^ with minor modifications. Briefly, starter cultures were grown overnight at 37 °C in LB medium supplemented with 0.3 M NaCl and under antibiotic selection. Cultures were diluted 1:10 in 50 ml of the same medium without antibiotics. The expression of *hilA* was induced by addition of 0.012% (w/v) L-arabinose for another growth period of 5 h to allow for the assembly of functional injectisomes and expression of effector proteins. Afterwards, the cell density was adjusted with LB media to an OD_600_ of 1.0 and the samples were centrifuged (6000 g for 15 min) to pellet the cells. Supernatants were collected and filtered with a 0.22 µM syringe filter to remove any residual cells. Pellets were resuspended in 1x PBS (phosphate buffered saline). Both supernatants and cell pellets were then immunoblotted with primary polyclonal rabbit antibodies raised against proteins of interest and secondary anti-rabbit HRP (horse radish peroxidase) conjugated antibodies (Qiagen).

Antibodies used in this study are summarized in Supplementary Table 6.

#### Needle complex purification

Preparation of the needle complexes was performed as described previously ^6, 7^. For assessing InvG and PrgH mutant/deletion versions, the protocol was scaled down by a factor of 4.

For selective needle complex disassembly, purified complexes were incubated with 20% v/v EtOH for 5 min at ambient temperature. The mixture was diluted with an equal volume of FR3 buffer (10 mM Tris-HСl, pH 8.0, 0.5 M NaCl, 5 mM EDTA., 0.1% l, pH 8.0, 0.5 M NaCl, 5 mM EDTA., 0.1% LDAO) and loaded onto a Superose 6 10/300 GL column equilibrated with FR3 buffer at 4°C. Fractions containing selectively disassembled needle complexes were pooled together, and analyzed by negative staining transmission electron microscopy. Optionally, samples were concentrated using centrifugal concentrators with a 30 kDa molecular weight cutoff.

#### Co-variation/Evolutionary coupling analysis

Co-variation between pairs of T3SS proteins was assessed using the EVCouplings complex pipeline ^8^ and RaptorX ^9^. For EVCouplings the results were filtered to have a coupling score above 0.05, and probability score above 0.8; results with discrete EVC ratio were not considered. For RaptorX results the probability cutoff was 0.5. All results are summarized in Supplementary Table 2.

#### Negative-stain electron microscopy

Carbon coated copper grids (400mesh) were glow discharged (Bal-Tec, SCD 005) for 40 seconds at 20 mA. Five µl of sample was applied to the grid and incubated for 30 seconds. The sample was washed off with 5 µl of staining solution (2% phospho-tungsten acetate, adjusted to pH 7.0 with NaOH) and stained with 5 µl of the staining solution for 20 sec. Micrographs were obtained at Morgagni 268D microscope (FEI) equipped with an 11 megapixel CCD camera (Olympus-SIS, Morada) or at a Talos L120C (FEI) with a 4K Ceta CEMOS camera.

#### Measurement and statistical analysis of inner ring diameters

Pixel intensity values corresponding to the inner ring were extracted from class average images and averaged along the vertical Y-axis for each class. IR diameters were calculated as the distance between the two minima of the averaged profile (Figure 1c,d, Supplementary Figure 1). Resulting diameter values were tested for normality using either D’Agostino-Pearson omnibus K2 test (for n=8 and above) or Shapiro-Wilk test (for n <8) with 95% confidence interval. Levene’s test with 95% confidence interval was used to assess the equality of variance. Unpaired t-test with 95% confidence interval was used to compare diameter measurements of mutants to the WT diameter. p-values are provided in Supplementary Table 1. Statistical analysis was performed in Python 2.7 using modules scipy.stats ^10^ and pandas ^11^.

#### Sample vitrification and cryo electron microscopy

For cryo electron microscopy, samples where vitrified on Quantifoil 1.2/1.3, 2/1, and 2/2 with either an additional home-made layer of amorphous carbon (<1.6nm) or graphene oxide ^12^. Briefly, 4ul of sample was applied onto glow-discharged grids and allowed to disperse for 0.5-2min. The grids were blotted for 4-7 s set at 100% humidity and plunge-frozen in liquid propane/ethane mixture cooled with liquid nitrogen to about minus 180-190 °C, by using a Vitrobot Mark V.

Vitrified specimens were imaged on a FEI Titan Krios operating at 300 kV and equipped with a field emission gun (XFEG) and a Gatan Bioquantum energy filter. Movies consisting of 25 frames were automatically recorded using FEI EPU software and the K2 Summit camera in counting mode at a nominal magnification of 130kx, corresponding to 1.09 Å per physical pixel. For individual frames, an electron dose of 1.1-1.26 e^-^/Å^2^ was used, corresponding to a cumulative electron dose of 27.5-31.5 e^-^/Å^2^ equally distributed over 5 sec movie. Movies were recorded at −0.8 - 3.5 µm defocus. Samples for diameter measurements were recorded with LEGINON^13^ on a FEI Polara (300kV) equipped with field emission gun (FEG) and a Gatan CCD Camera (UHS 4000). Total electron dose was 35 e^-^/Å^2^. The information on the datasets is summarized in Supplementary Table 3.

#### Single-particle reconstruction

Single particle reconstructions were performed using Relion3^14^. Movies were motion-corrected and dose-weighted and the CTF of the resulting micrographs was determined using CTFFIND4 ^15^. Particles were picked from the motion-corrected micrographs using Cryolo ^16^ trained with a sub-set of manually picked particles (4-fold binned. Particles were extracted using a boxsize of 432 pixels and subsequently binned four times for several rounds of 2D classifications. A cleaned and unbinned dataset was obtained by re-extraction and aligned to a rotationally averaged structure. Focused refinements with and without applying symmetry were preformed to the individual sub-structures (Supplementary Table 4) using respective 3D masks. After converged refinements, per-particle CTF and Bayesian polishing was used to generate new data sets for another round of focused refinements. Overall gold-standard resolution (Fourier shell correlation (FSC) =0.143) and local resolution were calculated with Relion3. (Data sets from WT needle complex recorded over amorphous carbon and graphene oxide were processed separately and combined into a single data set after Bayesian polishing).

#### Helical reconstruction of the outer needle filament

Helical reconstruction was performed in Relion3 ^17^ on the carbon dataset used for high-resolution refinement of the needle complex. The box center was moved from its original position at the basal body to the approximate middle of the filament by recalculating the particle coordinates according to the formulae:

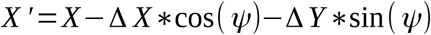

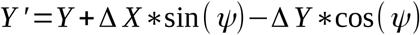

Where X, Y are the initial particle coordinates (rlnCoordinateX and rlnCoordinateY fields in *.star file), ΔX and ΔY are the shifts of the particle as measured on a 2D class average and ψ is the in-plane rotation angle (rlnAnglePsi field in *.star file).

Re-extracted particles were subjected to 2D classification. Classes where the helicity of the filament was most pronounced were used to convert box center coordinates to helix start and end coordinates using the formula above. Calculations of new coordinates were done with Python 2.7 and modules pandas and numpy ^11, 18^. Segments were extracted from micrographs into 300 pixel boxes with 28 pixel inter-box distance (defined by helical rise of 4.2 Å and 6 asymmetric units). In total 303 155 segments were extracted. The dataset was classified in 2D with tube diameter set to 95 Å and fine angular sampling. Classes that showed no evidence of overfitting and little or no extra density from the IR were selected for 3D auto-refinement procedure (total 133516 segments). Tube diameter was set to 95 Å, and the initial values for helical twist and rise were 64° and 4.2 Å (as determined in ^19^). Helical parameter search range was ± 20%, and a soft-edged cylinder with diameter 95 Å was used as an initial model. The resulting model was then subjected to CTF refinement and Bayesian polishing. Where applicable the initial models were low-pass filtered to 25 Å to keep the helical lattice intact while removing any high-resolution components. The map, filtered according to its local resolution in Relion, was symmetrized in real space using the refined values of helical twist and rise to generate the final map.

#### Building, refinement and validation of needle complex models

##### Templates and initial model generation

Existing models of PrgH (PDB: 3GR1) and PrgK (PDB: 3GR5), and InvG (4G08, G34-I173; lower OR) were preliminary placed into the respective EM maps (IR: WT 24-mer, ΔEA 23-mer; lower OR: WT 16mer, ΔEA 15mer; Supplementary Table 4) utilizing UCSF Chimera’s `fitmap` command ^20^. The models were further refined using Rosetta ^21^ controlled via StarMap 1.0 (manuscript in preparation).

The upper OR was modelled into the 15mer symmetrized EM map by building InvG (R177-G557) ab initio with Coot^22^, starting with well defined secondary structure elements. The growing model was iteratively refined using Rosetta (controlled by Starmap), followed by several rounds of model extension and manual improvement in Coot.

Models for the upper and lower OR were connected in Coot based on densities obtained through focused refinements of the neck region.

Homology models for SpaP, SpaQ, SpaR, SpaS were generated with SWISS-MODEL ^23^ based on FliP, FliQ, FliR, and FliS from Vibrio mimicus (personal communication, S. Lea); PrgJ was modelled using Phyre2 ^24^. PrgI (PDB 2LPZ) and models for SpaP, SpaQ, SpaR, SpaS, and PrgJ were preliminary placed into the respective EM map, extended if necessary, and refined using Rosetta (Starmap) and Coot.

##### Model refinement and validation

All resulting models were manually extended and refined using Coot, and subsequently subjected to another round of refinement using Rosetta. The quality of the individual models were analyzed using the MolProbity server ^25^ and parts of the models further refined if necessary.

Overall map/model FSCs as well as Z-scores ^26^ were calculated with Starmap to assess the quality of the models. Z-scores were colored for each model using Chimera ^20^.

#### Visualization, analysis and deposition

UCSF Chimera ^20^ and ChimeraX ^27^ were used for molecular visualization. Analysis of interaction surfaces between inner ring components PrgH and PrgK in the WT and ΔEA T3SS needle complex were performed with the PDBePISA server ^28^. Angles in InvG were measured with MB-Ruler (MB-Software solutions, version 5.3). Plots were made with matplotlib ^29^.

EM maps are deposited at the EMDB (https://www.emdataresource.org). Coordinates of models are deposited at the ePDB database. Supplementary Table 4 summarizes the depositions submitted in this study.

## Supplementary figures

**Supplementary Figure 1:**
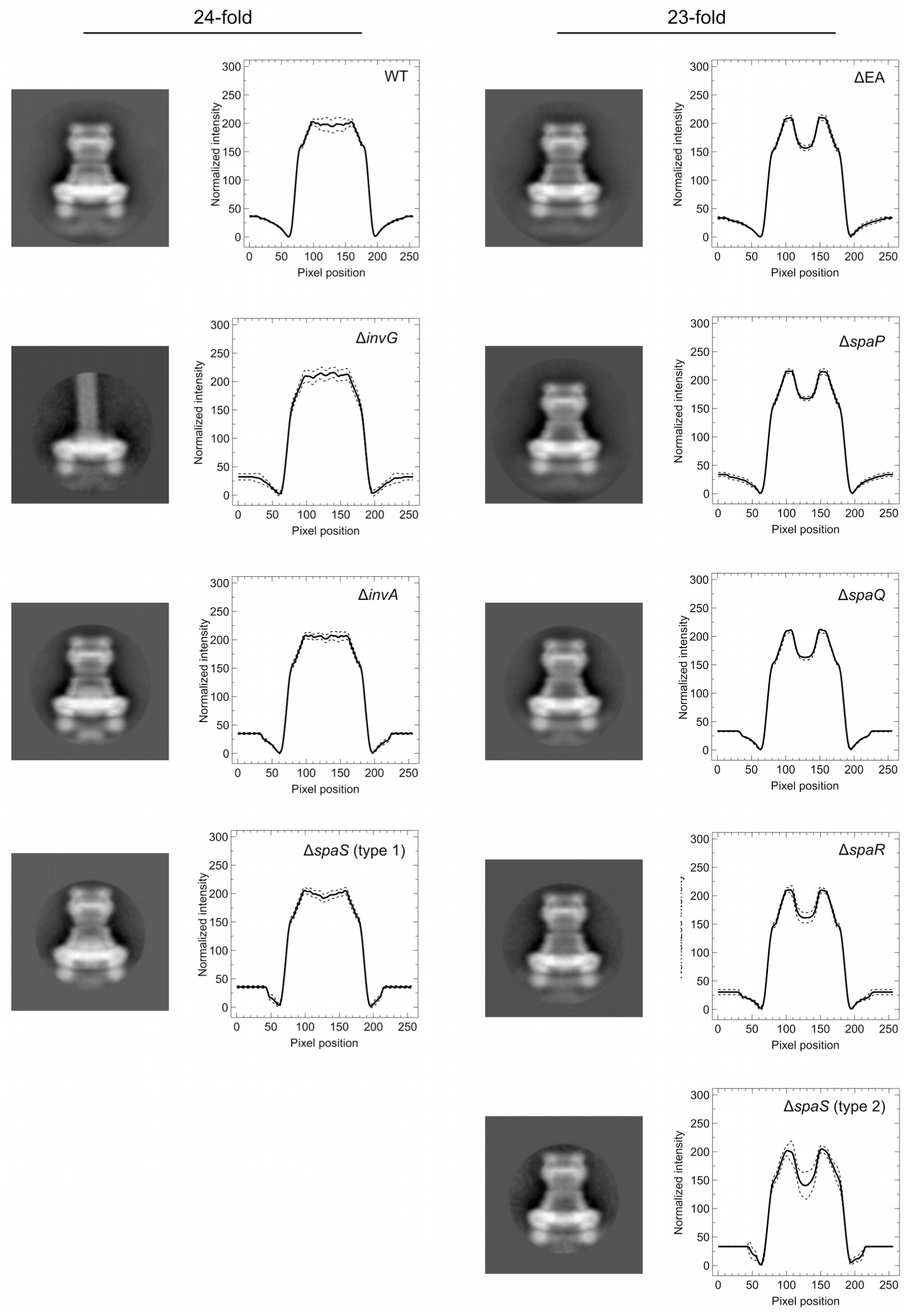
T3SS-complexes from WT and mutant strains display different IR diameters. Two-dimensional class averages from vitrified complexes (WT, ΔEA, Δ*invG*, Δ*invA,* Δ*spaS,* Δ*spaP,* Δ*spaQ,* Δ*spaR)* and density profile along IR. Diameters were measured as the distance of the two lowest minima along the density profile (dashed lines: standard deviation) and summarized in Supplementary Table 1.

**Supplementary Figure 2:**
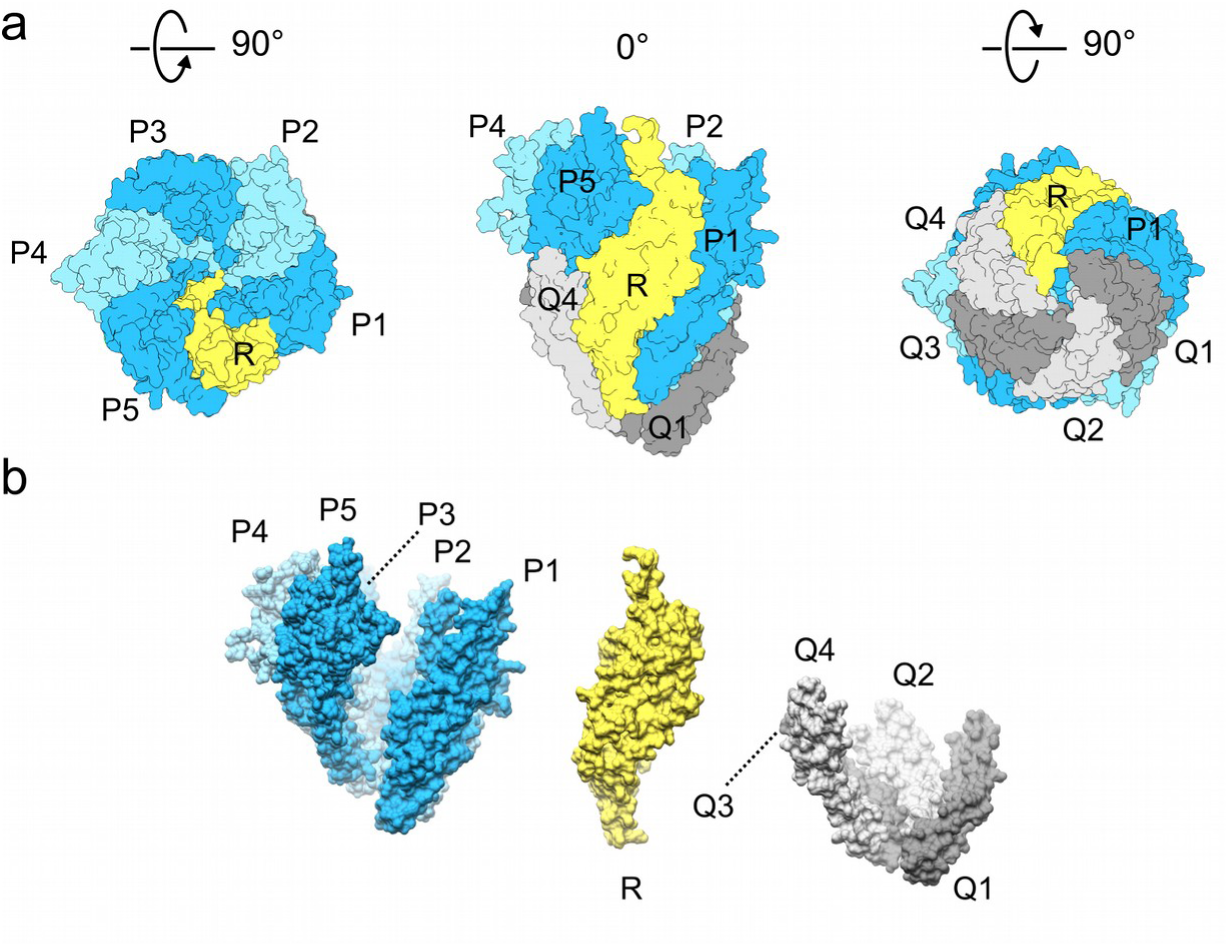
The PQR complex adopts a pseudo-helical assembly. a) Surface views from the PQR complex at different orientations (top, side, bottom). Proteins are colored according to their identity (SpaP: blue; SpaR: yellow, SpaQ: grey) b) Surface views of the pseudo-helical packing of the individual SpaP-group, SpaR, and SpaQ-group. SpaQ resembles the bottom of the PQR complex, whereas the SpaP group is arranged at the top-rim of the complex

**Supplementary Figure 3:**
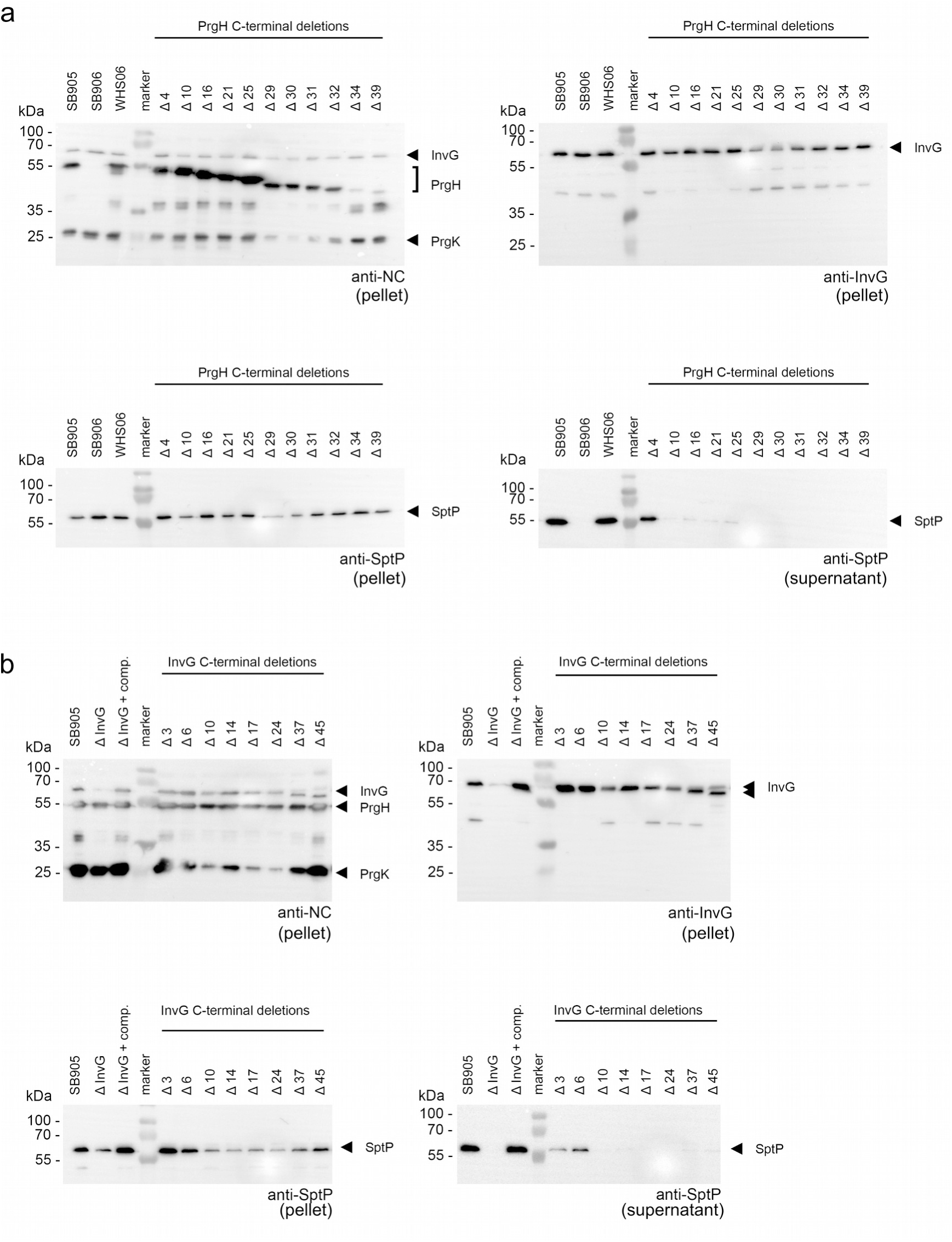
C-terminal truncations of PrgH or InvG impair T3SS function. a) Secretion assays for C-terminal PrgH truncations. Deletions of four amino acids do not, and more than 10 amino acids render the T3SS non-functional. Left/top: Detection of needle complex components (NC) in cell pellets, Top/right: Detection of InvG in PrgH-truncated cells; Bottom/left: Detection of late T3SS-substrate SptP in cell pellets. Bottom/ right: Detection of secreted late T3SS-substrate SptP in cell culture supernatants. (WT: SB905, SB906: Δ*prgH*, WHS06: Δ*prgH* (SB906), complemented with plasmid-borne WT *prgH*), Δ4-Δ39: Δ*prgH* (SB906) complemented with plasmid-borne C-terminal PrgH deletions; anti-NC (anti needle complex antibody). b) Secretion assasy for C-terminal InvG truncations. Deletions of six amino acids do not, and more than 10 amino acids render T3SS non-functional. Left/top: Detection of needle complex components (NC) in cell pellets, Top/right: Detection of InvG and truncated InvG in cells; Bottom/left: Detection of late T3SS-substrate SptP in cell pellets. Bottom/right: Detection of secreted late T3SS-substrate SptP in cell culture supernatants. (WT: SB905, Δ*invG*: SB908, Δ*InvG* + comp.: SB908 complemented with plasmid-borne WT *invG*, Δ3-Δ45: Δ*invG* complemented with plasmid-borne C-terminal InvG deletions; anti-NC (anti needle complex antibody).

**Supplementary Figure 4:**
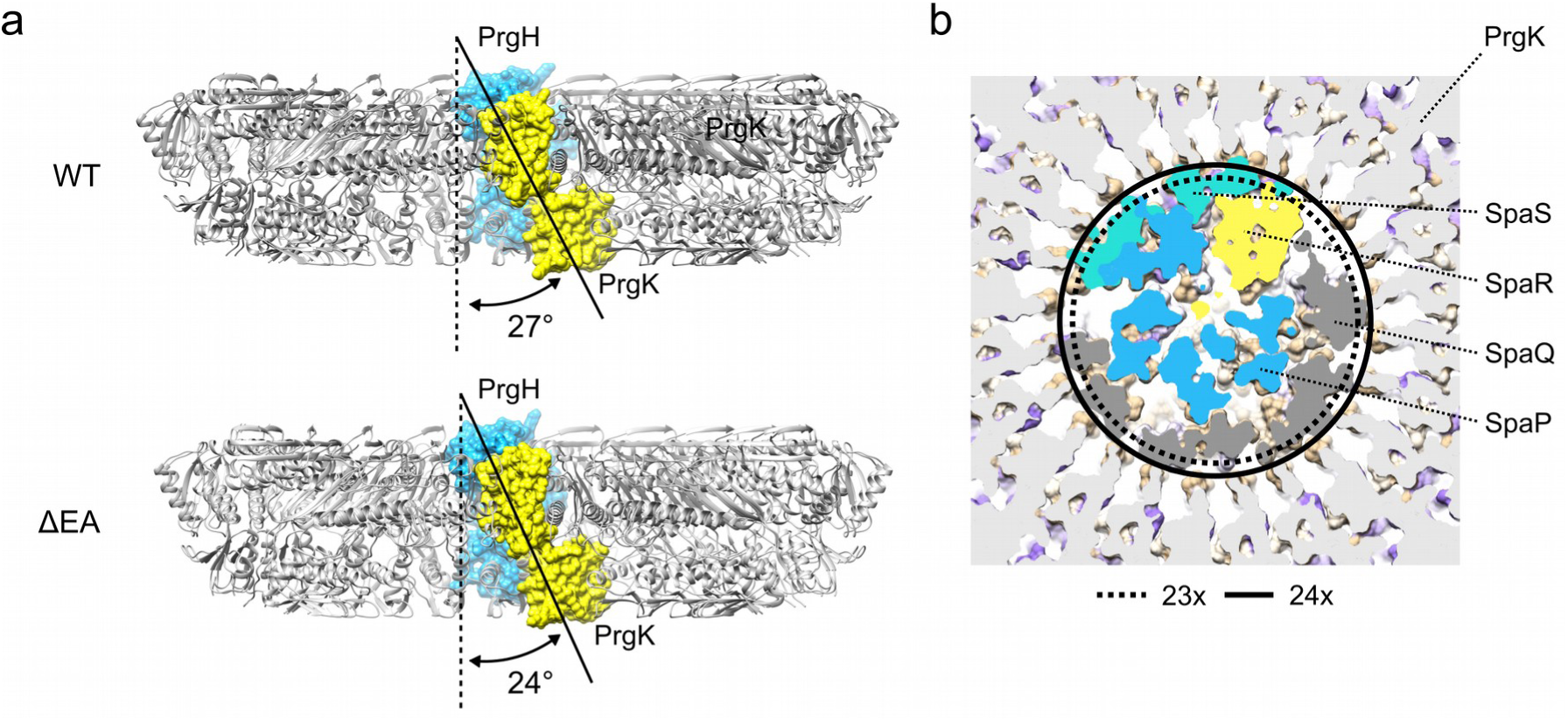
The dead-end injectisome assembly. a) PrgH and PrgK engage in a similar intertwined packing to from the IR in WT and ΔEA complexes. PrgH (blue, one protomer) is oriented perpendicular relative to the IR, whereas PrgK (yellow, one protomer) binds to the PrgH-ring at an angle between 24 (ΔEA) and 27°. (WT). In this view, the upper PrgK domain binds to PrgH shown in blue; the lower PrgK domain binds to the neighboring PrgH protomer (+1). b) The PQRS complex does not fit into a 23-mer complex, and provides a structural template for the WT 24-mer IR. Inner circle defined by the 24-mer WT complex (continuous line) and the 23-mer ΔEA complex (dashed line). Surface view through IR, SpaP (blue), SpaR (yellow), SpaQ (dark grey), SpaS (green). The surface of PrgK is colored according to hydrophobicity (see also Figure 2), and the section depicted in light grey.

**Supplementary Figure 5:**
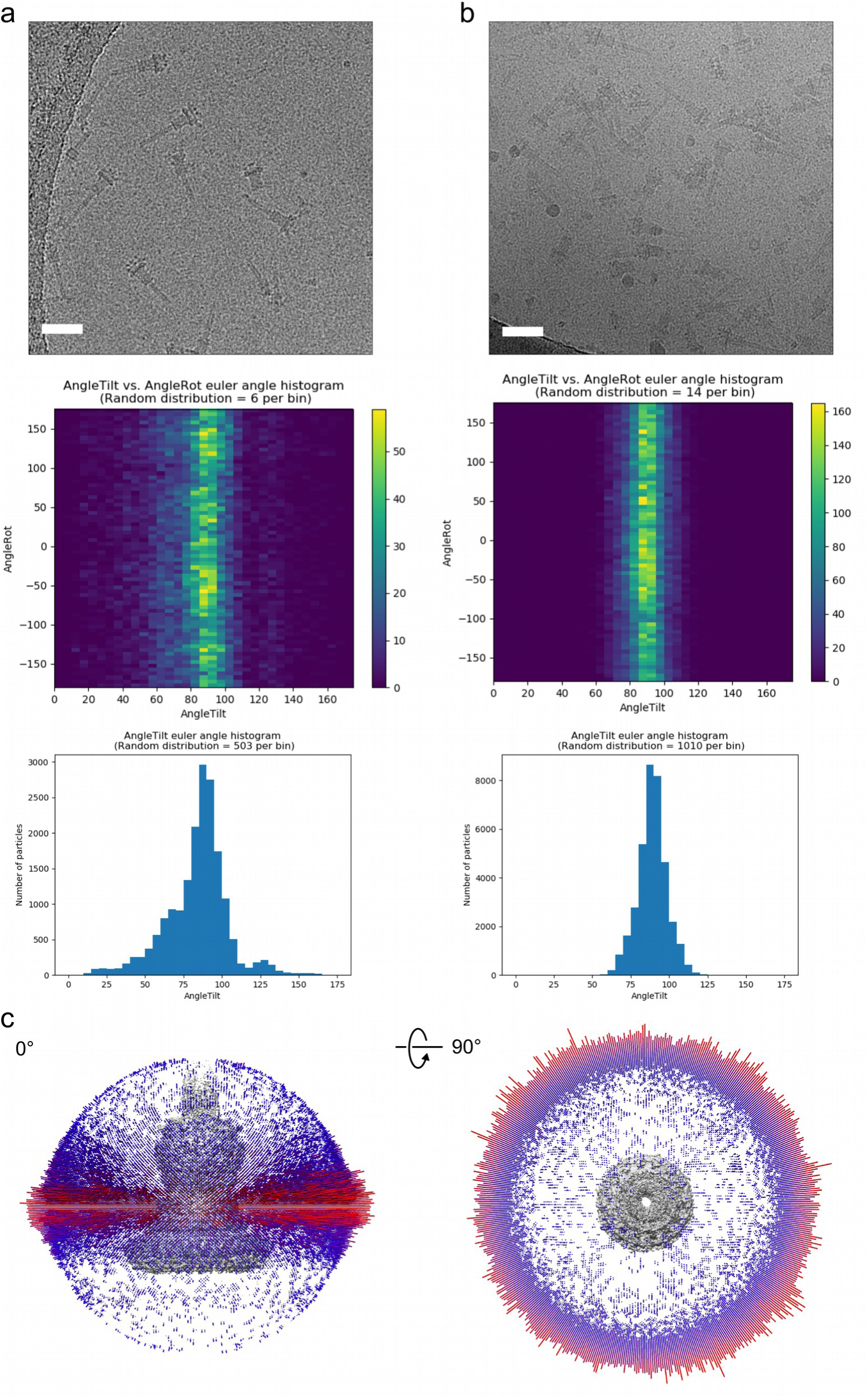
Grid support type affects needle complex particle distribution in cryo electron microscopy. a) Exemplary data for needle complexes vitrified on grids containing a graphene oxide support layer. Top: Cryo electron micrograph (5-fold binned and 25 Å low-pass filtered in Relion), scale bar 50 nm. Middle: Histograms of Euler angle distribution (Rot and Tilt) of asymmetric 3D refinement created using the plot_indivEuler_histogram_fromStarFile.py (Michael A. Cianfrocco, U. Michigan) and histogram of Euler angle distribution (Tilt). b) Same as in (a), yet for continuous carbon grids. c) Angular distribution of C1 reconstruction (job373) of the sub-dataset focusing on the WT export apparatus complex. Distribution shown from the side and rotated 90° along the x-axis. Lighter colors (magenta) indicate a higher number of particles found with the respective tilt angle.

**Supplementary Figure 6:**
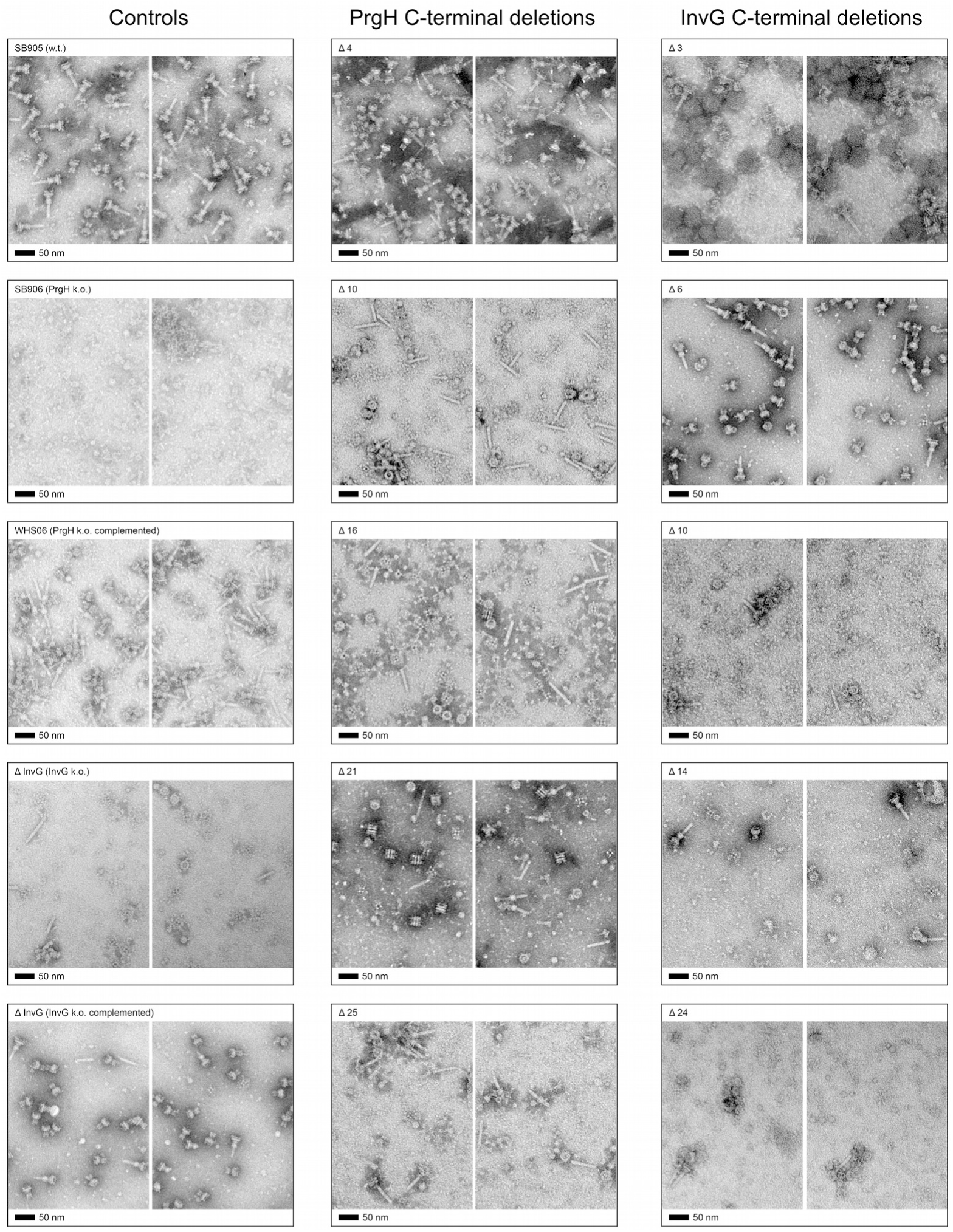
Stability of T3SS needle complexes is impaired by C-terminal PrgH or InvG truncations. Representative images of isolated complexes from WT, *ΔprgH*, *ΔinvG* and plasmid-complemented strains. Plasmid-borne truncated versions of PrgH and InvG are indicated. (Middle) PrgH C-terminal deletions – Δ4, 10, 16, 21 and 25; (Right) InvG C-terminal deletions – Δ3, 6, 10, 14 and 24; Scale bar 50 nm.

**Supplementary Figure 7:**
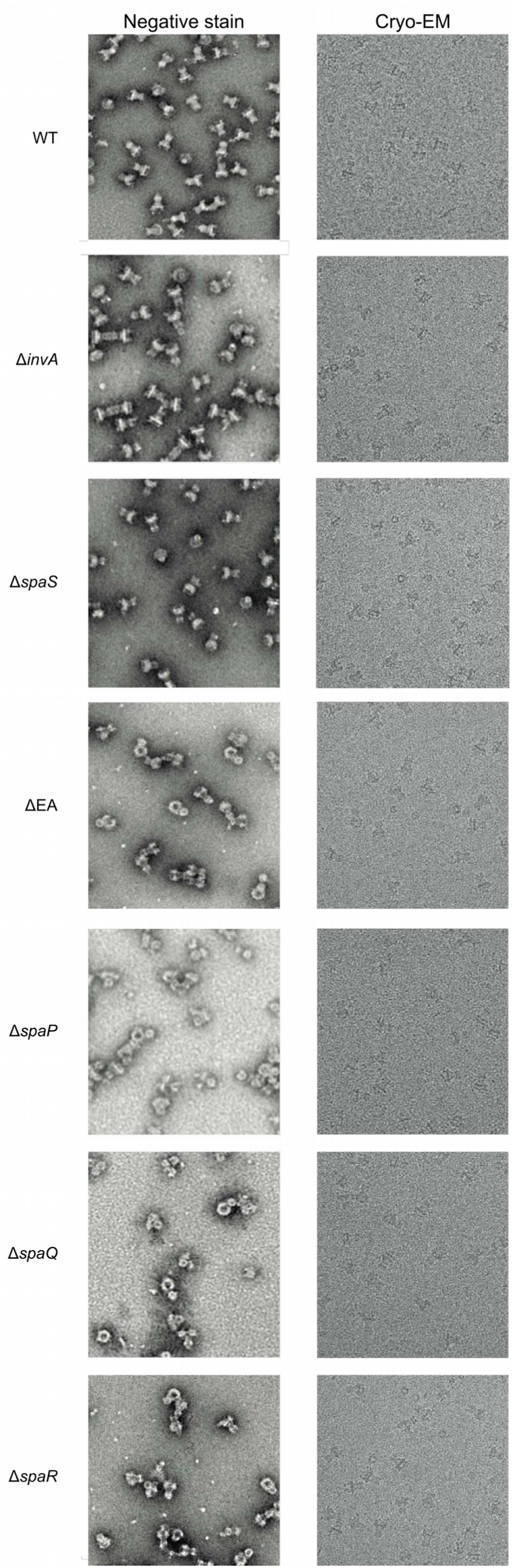
Export apparatus knock-out basal-bodies are unstable in negative-stain EM. Isolated complexes from a strain lacking the entire export apparatus proteins (ΔEA: Δ(*spaP*, *spaQ*, *spaR*, *spaS*, *invA*)), and the individual genes for *spaP, spaQ, spaR* are sensitive to structural disruption upon negative stain electron microscopy, likely through shear forces during the staining procedure. However, the same samples withstand vitrification for cryo electron microscopy.

**Supplementary Figure 8:**
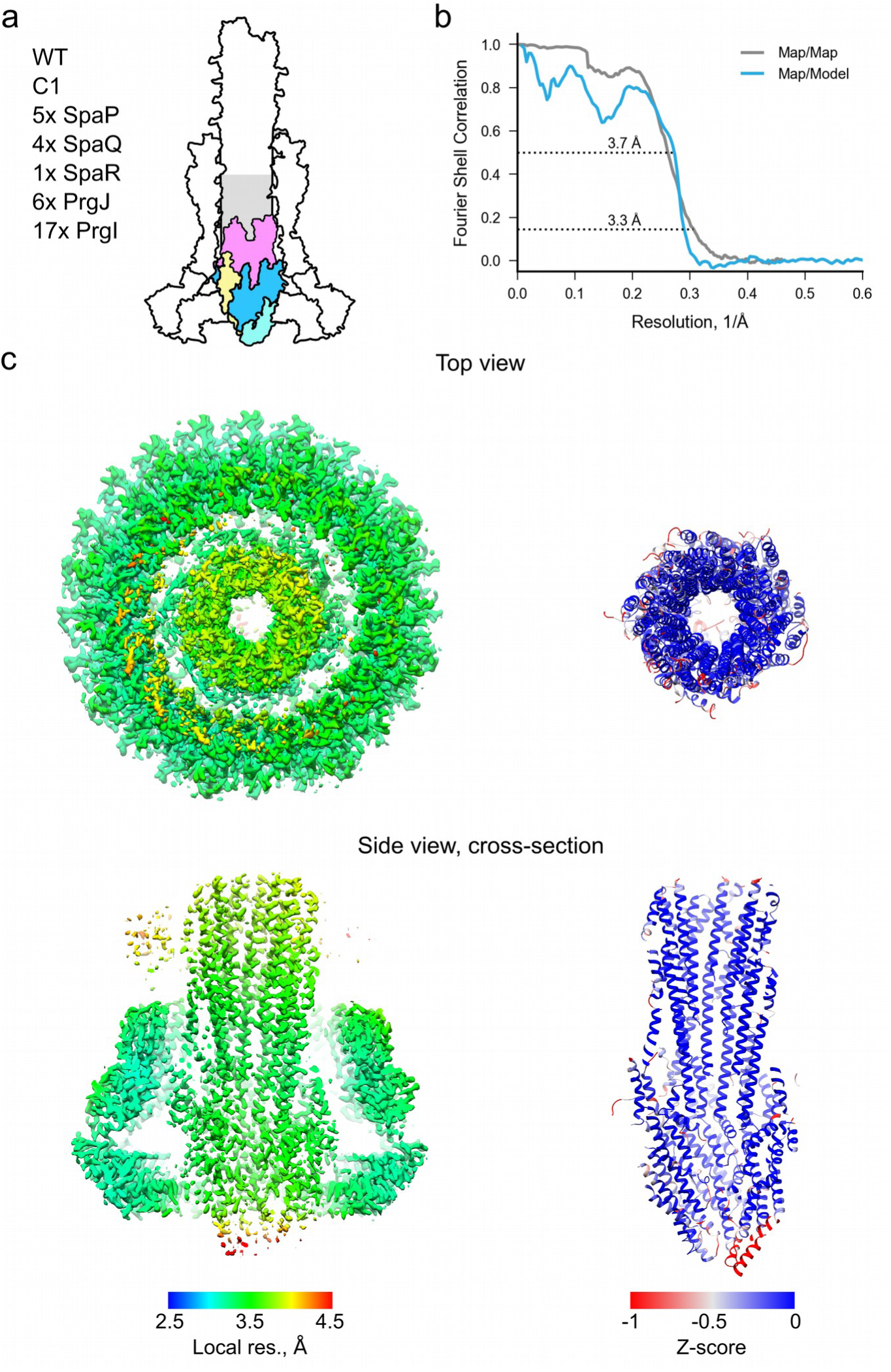
Summary for asymmetric reconstruction of WT export apparatus. a) Schematic showing the location of the complex in inside the needle complex b) Fourier-shell correlation plots of the: map versus map (grey) and map versus model (blue). The resolution at the respective cut-off is given. c) Left: post-processed electron density map, colored by local resolution (Relion implementation); right: ribbon view of the model colored by Rosetta Z-score

**Supplementary Figure 9:**
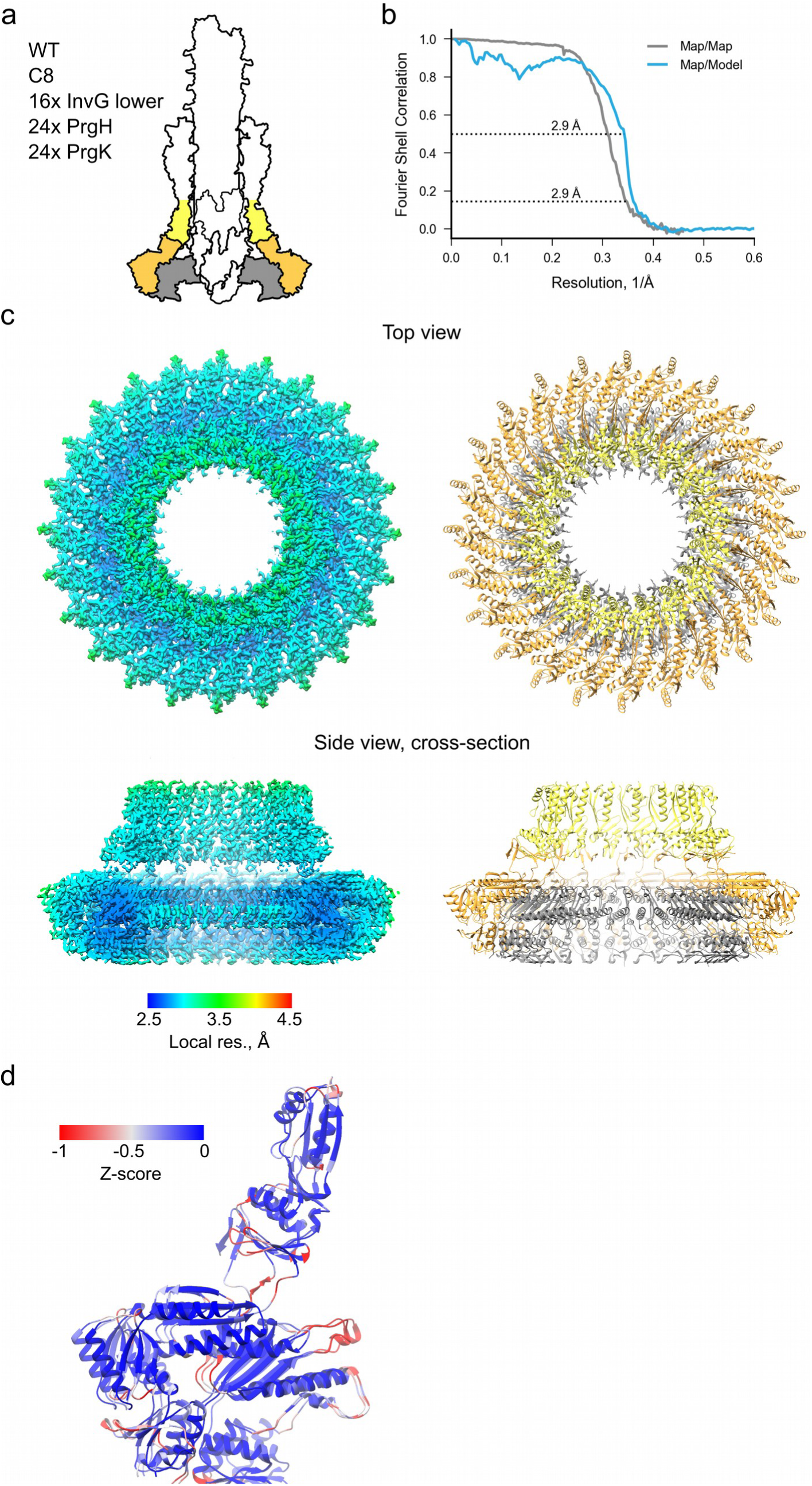
Summary for C8 reconstruction of WT IR and OR lower region. a) Schematic showing the location of the complex in inside the needle complex b) Fourier-shell correlation plots of the: map versus map (grey) and map versus model (blue). The resolution at the respective cut-off is given. c) Left: post-processed electron density map, colored by local resolution (Relion implementation); right: ribbon view of the model d) An asymmetric unit in ribbon view colored by Rosetta Z-score

**Supplementary Figure 10:**
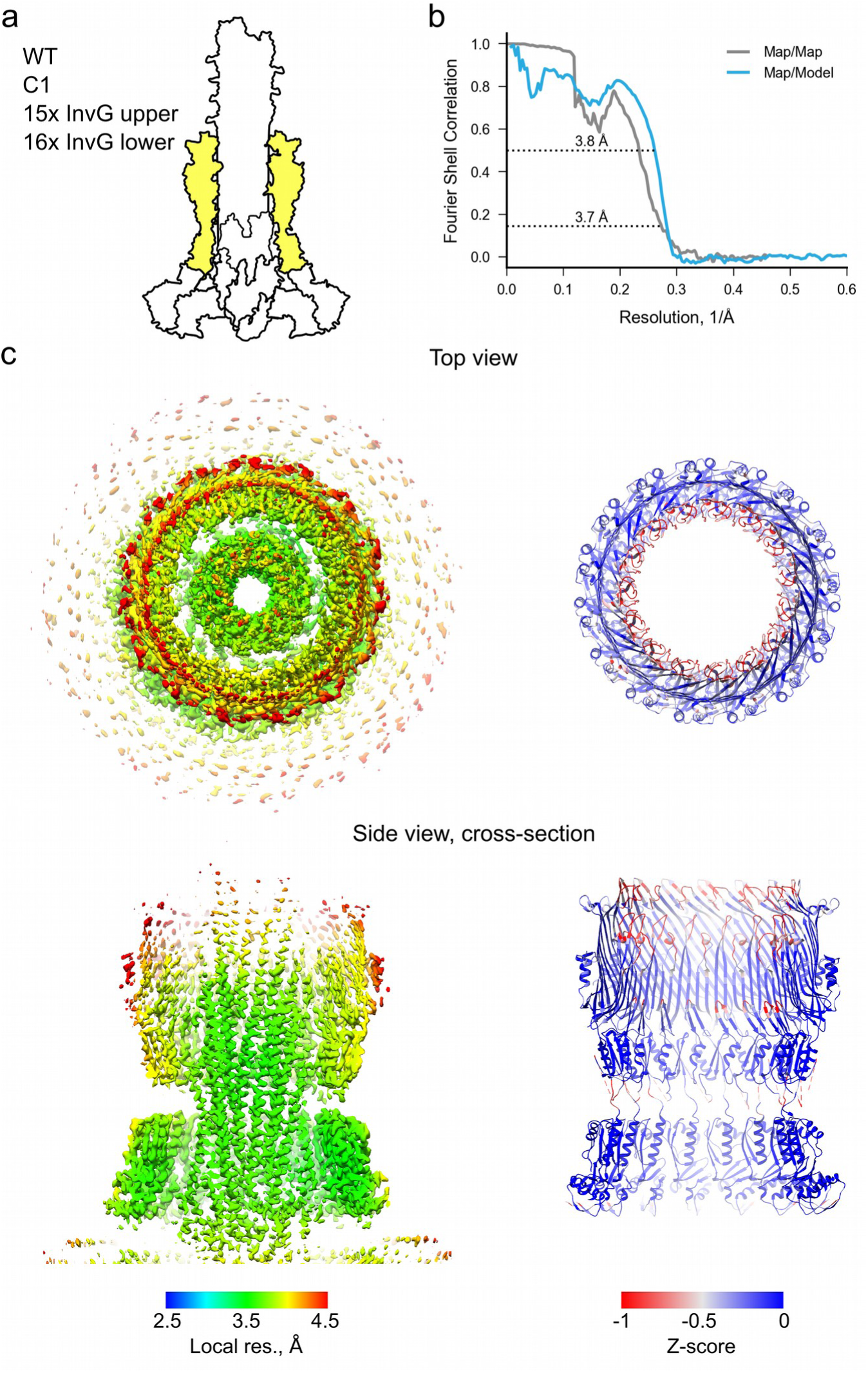
Summary for asymmetric reconstruction of WT OR. a) Schematic showing the location of the complex in inside the needle complex b) Fourier-shell correlation plots of the: map versus map (grey) and map versus model (blue). The resolution at the respective cut-off is given. c) Left: post-processed electron density map, colored by local resolution (Relion implementation); right: ribbon view of the model colored by Rosetta Z-score

**Supplementary Figure 11:**
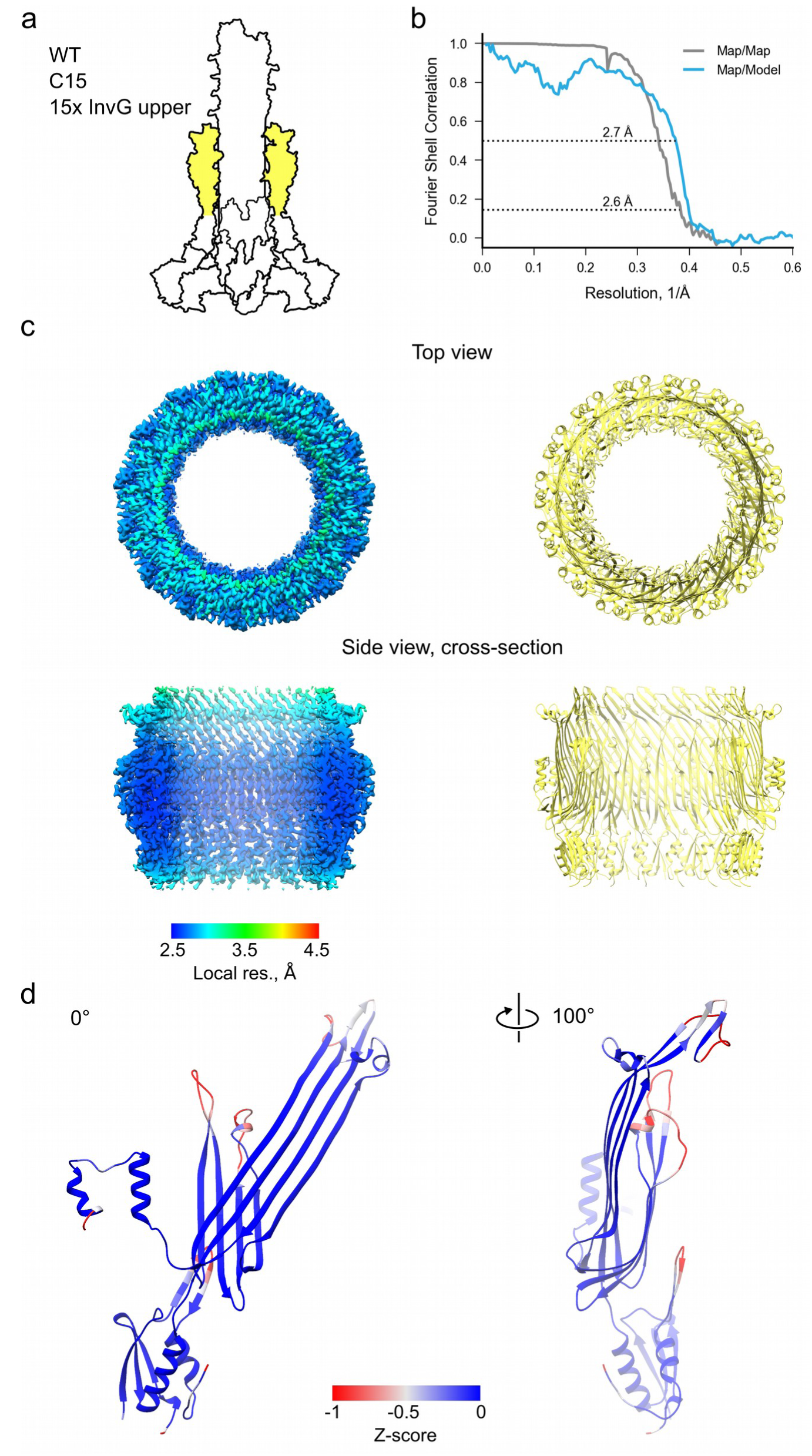
Summary for C15 reconstruction of WT InvG upper region. a) Schematic showing the location of the complex in inside the needle complex b) Fourier-shell correlation plots of the: map versus map (grey) and map versus model (blue). The resolution at the respective cut-off is given. c) Left: post-processed electron density map, colored by local resolution (Relion implementation); right: ribbon view of the model d) An asymmetric unit in ribbon view colored by Rosetta Z-score

**Supplementary Figure 12:**
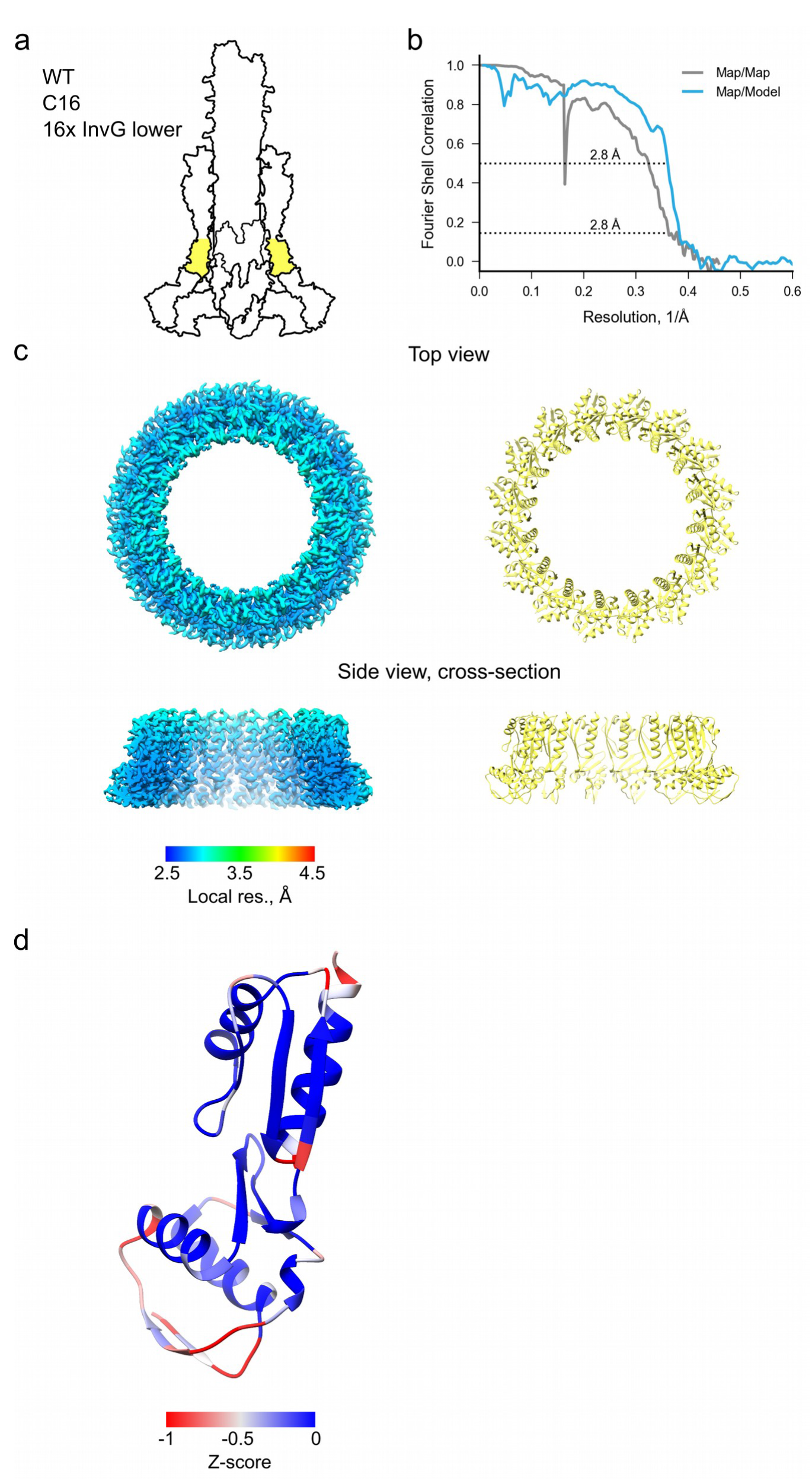
Summary for C16 reconstruction of WT InvG lower region. a) Schematic showing the location of the complex in inside the needle complex b) Fourier-shell correlation plots of the: map versus map (grey) and map versus model (blue). The resolution at the respective cut-off is given. c) Left: post-processed electron density map, colored by local resolution (Relion implementation); right: ribbon view of the model d) An asymmetric unit in ribbon view colored by Rosetta Z-score

**Supplementary Figure 13:**
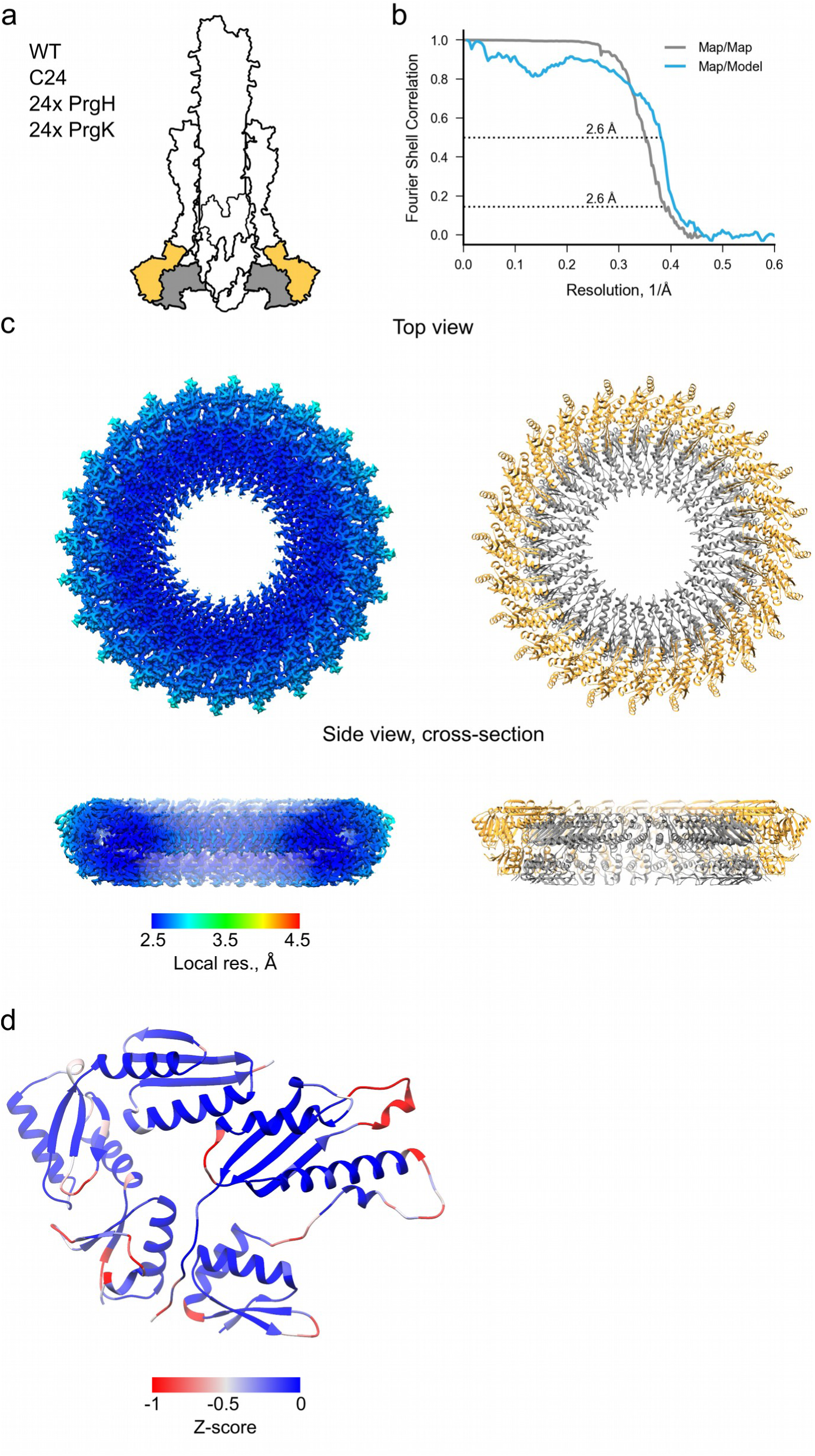
Summary for C24 reconstruction of WT IR. a) Schematic showing the location of the complex in inside the needle complex b) Fourier-shell correlation plots of the: map versus map (grey) and map versus model (blue). The resolution at the respective cut-off is given. c) Left: post-processed electron density map, colored by local resolution (Relion implementation); right: ribbon view of the model d) An asymmetric unit in ribbon view colored by Rosetta Z-score

**Supplementary Figure 14:**
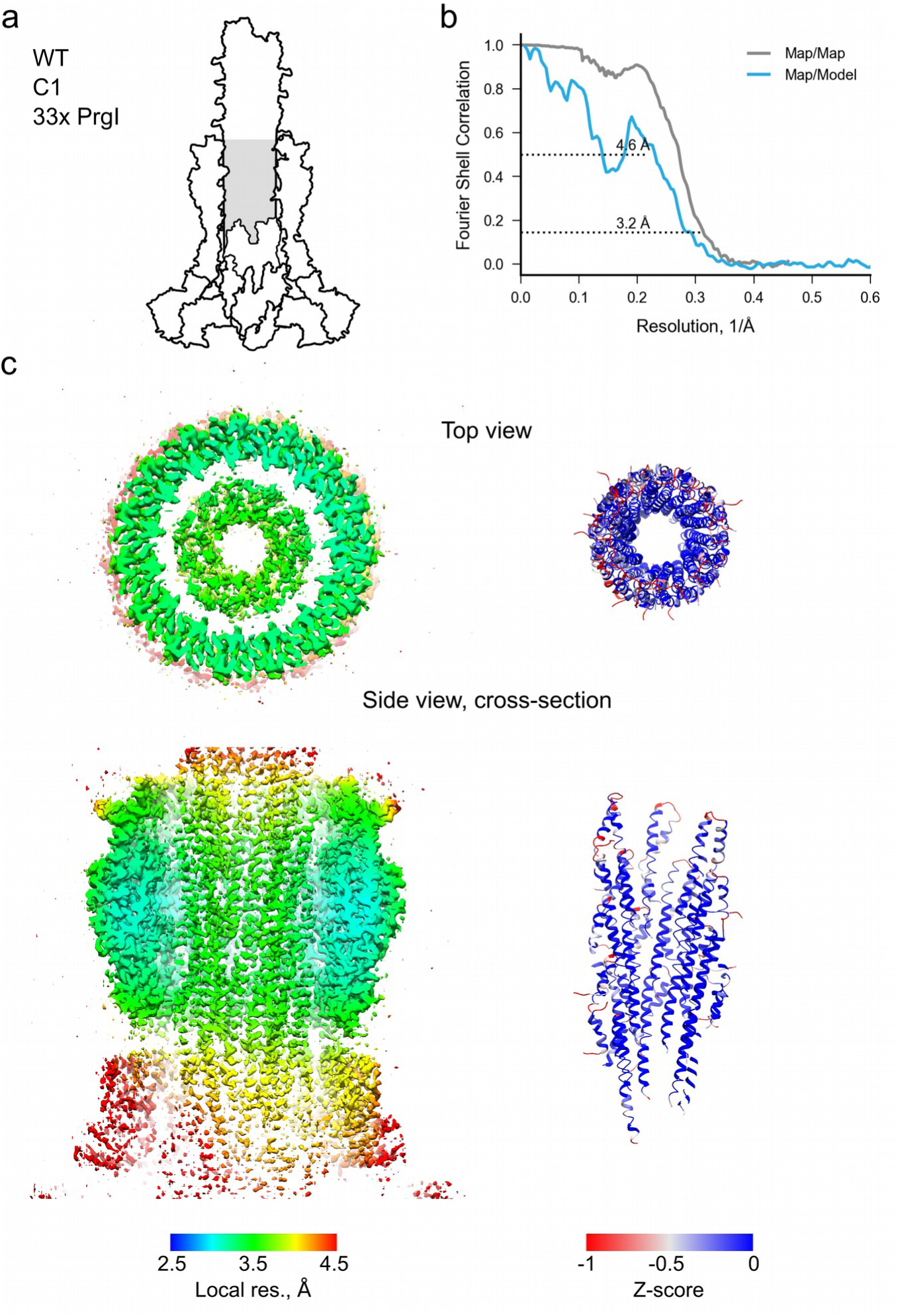
Summary for asymmetric reconstruction of WT inner filament part. a) Schematic showing the location of the complex in inside the needle complex b) Fourier-shell correlation plots of the: map versus map (grey) and map versus model (blue). The resolution at the respective cut-off is given. c) Left: post-processed electron density map, colored by local resolution (Relion implementation); right: ribbon view of the model colored by Rosetta Z-score

**Supplementary Figure 15:**
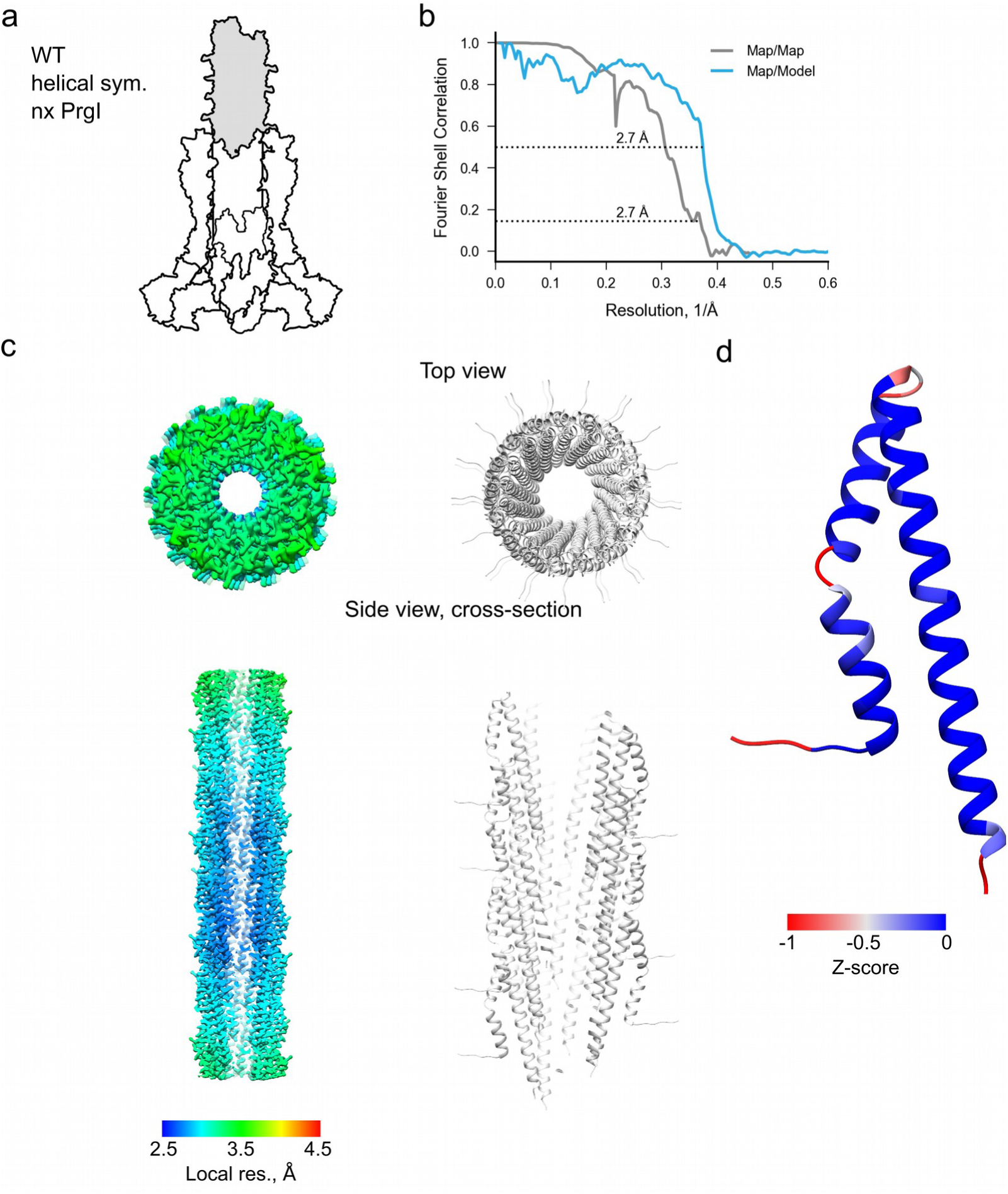
Summary for helical reconstruction of WT outer filament part. a) Schematic showing the location of the complex in inside the needle complex b) Fourier-shell correlation plots of the: map versus map (grey) and map versus model (blue). The resolution at the respective cut-off is given. c) Left: post-processed electron density map, colored by local resolution (Relion implementation); right: ribbon view of the model d) An asymmetric unit in ribbon view colored by Rosetta Z-score

**Supplementary Figure 16:**
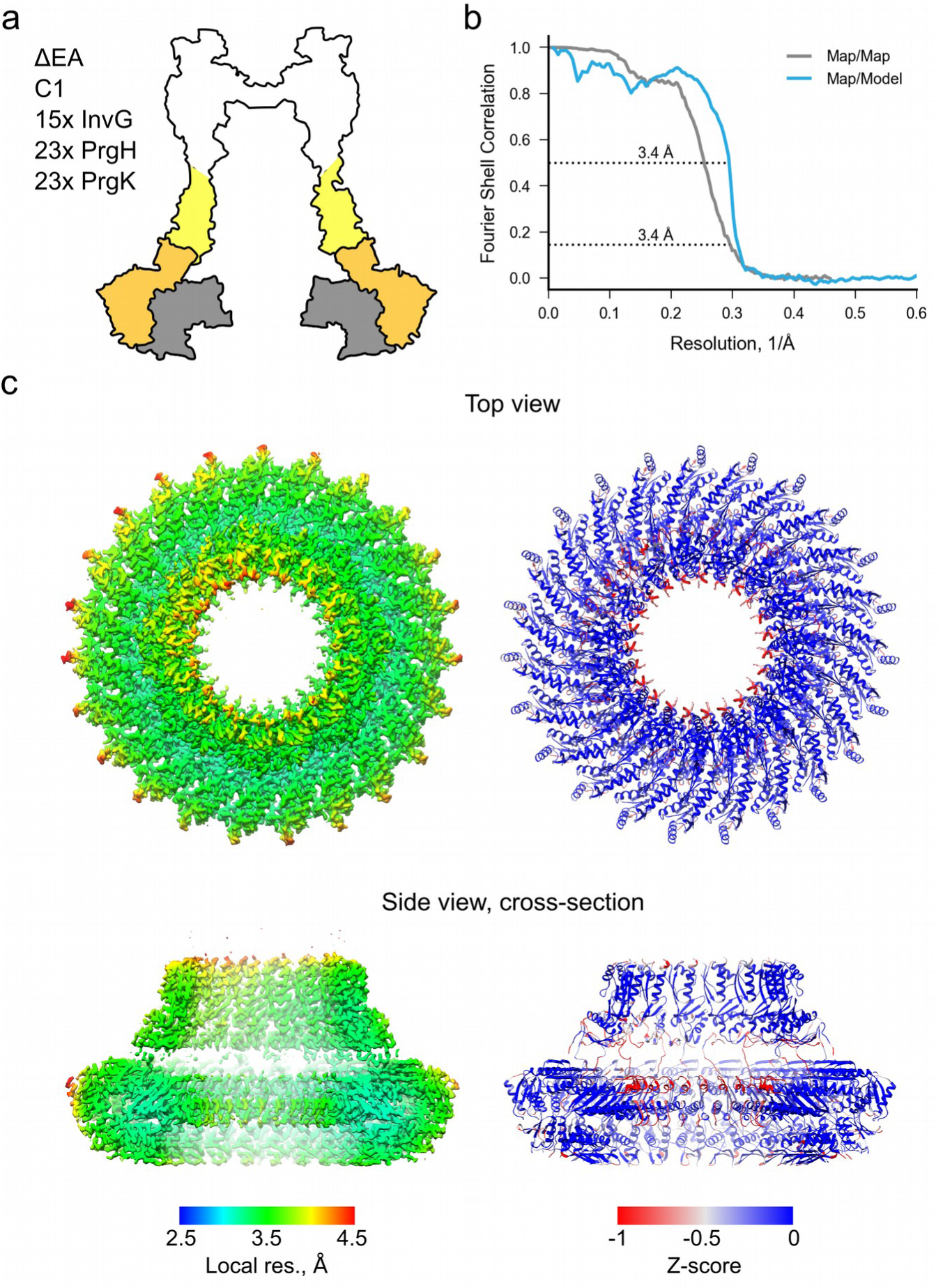
Summary for asymmetric reconstruction of ΔEA export EA export IR and InvG lower region. a) Schematic showing the location of the complex in inside the needle complex b) Fourier-shell correlation plots of the: map versus map (grey) and map versus model (blue). The resolution at the respective cut-off is given. c) Left: post-processed electron density map, colored by local resolution (Relion implementation); right: ribbon view of the model colored by Rosetta Z-score

**Supplementary Figure 17:**
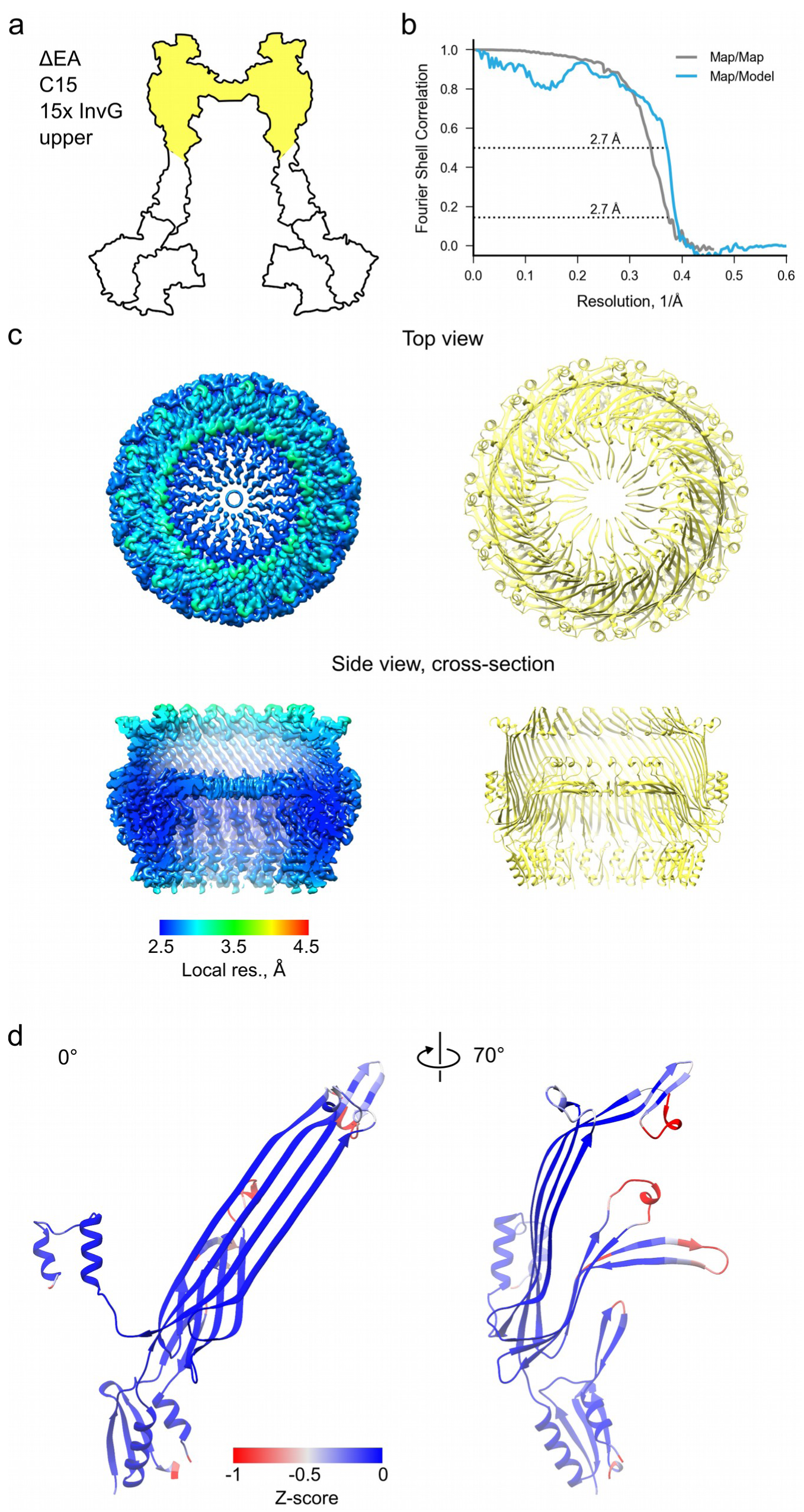
Summary for C15 reconstruction of ΔEA export EA InvG upper region. a) Schematic showing the location of the complex in inside the needle complex b) Fourier-shell correlation plots of the: map versus map (grey) and map versus model (blue). The resolution at the respective cut-off is given. c) Left: post-processed electron density map, colored by local resolution (Relion implementation); right: ribbon view of the model d) An asymmetric unit in ribbon view colored by Rosetta Z-score

**Supplementary Figure 18:**
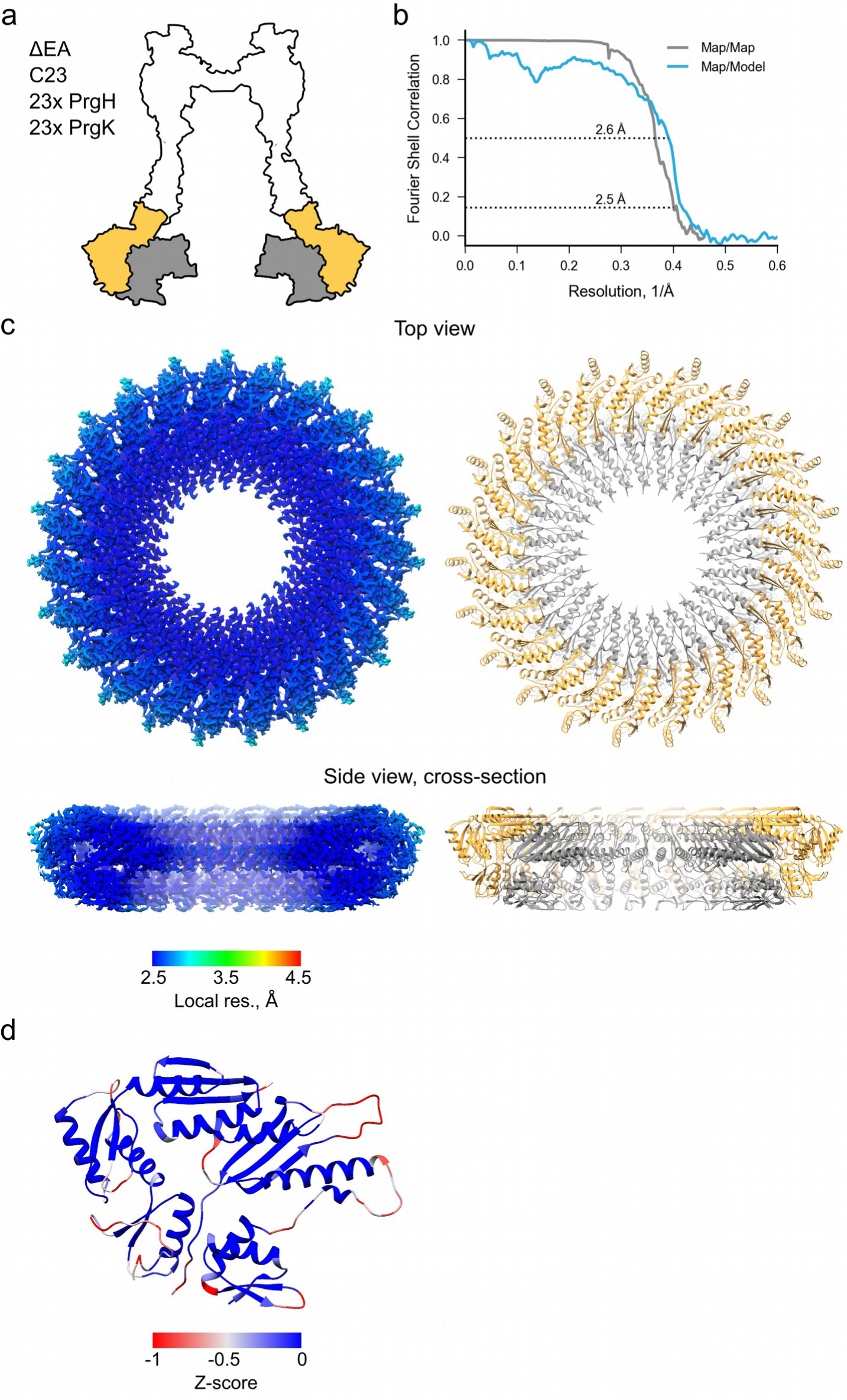
Summary for C23 reconstruction of ΔEA export EA IR. a) Schematic showing the location of the complex in inside the needle complex b) Fourier-shell correlation plots of the: map versus map (grey) and map versus model (blue). The resolution at the respective cut-off is given. c) Left: post-processed electron density map, colored by local resolution (Relion implementation); right: ribbon view of the model d) An asymmetric unit in ribbon view colored by Rosetta Z-score

**Supplementary Figure 19:**
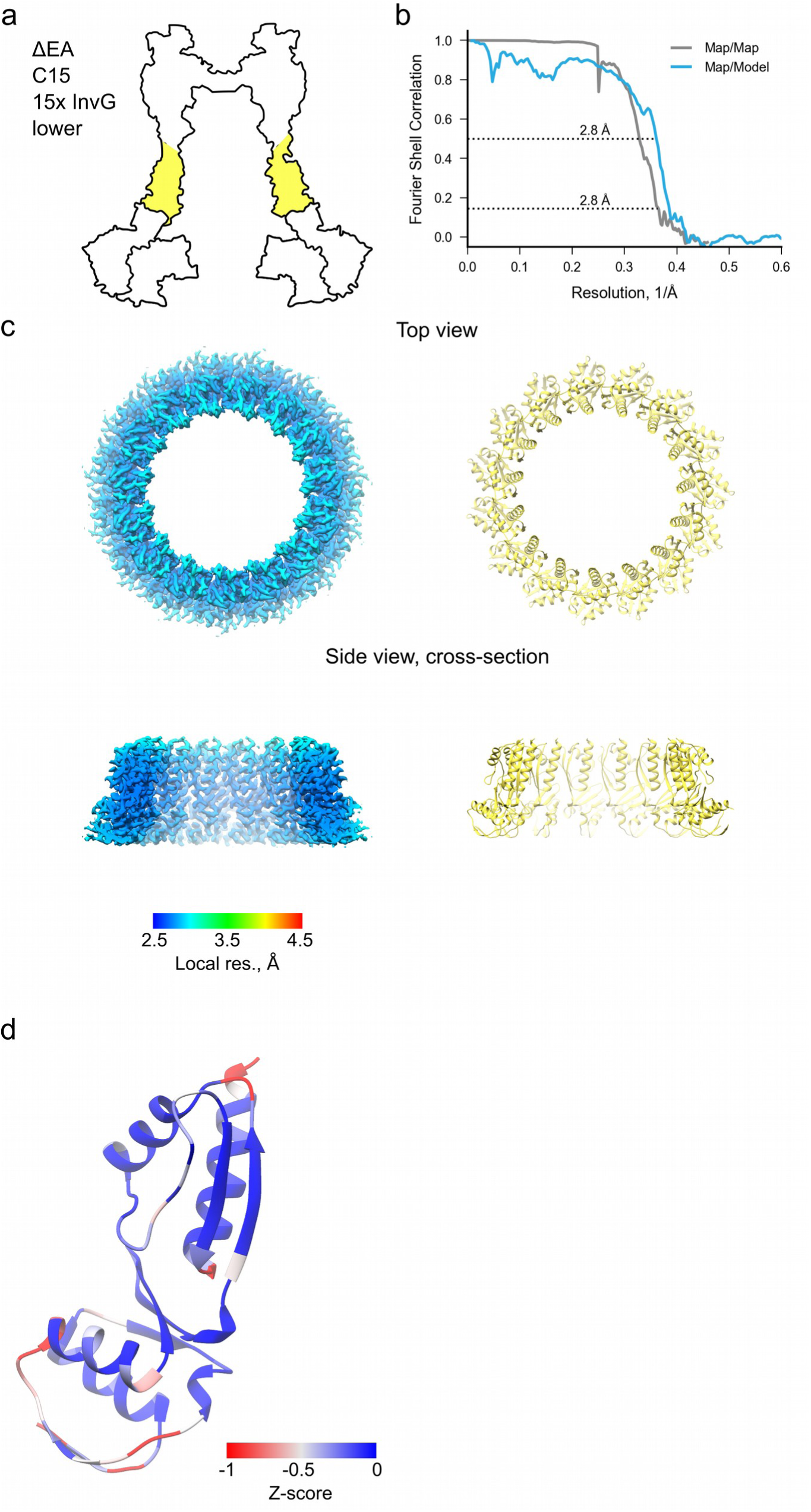
Summary for C15 reconstruction of ΔEA export EA InvG lower region. a) Schematic showing the location of the complex in inside the needle complex b) Fourier-shell correlation plots of the: map versus map (grey) and map versus model (blue). The resolution at the respective cut-off is given. c) Left: post-processed electron density map, colored by local resolution (Relion implementation); right: ribbon view of the model d) An asymmetric unit in ribbon view colored by Rosetta Z-score

**Supplementary Table 1:** Statistical analysis of IR diameters from 2D class averages.

| Sample | n | Diameter, pix | Normality test | Normality test p-value | Normal distribution | Levene test p-value | Equal variance to WT | t-test p-value | IR oligomericity |
| --- | --- | --- | --- | --- | --- | --- | --- | --- | --- |
| WT | 8 | 135.88±0.99 | D'Agostino-Pearson | 0.03 | No | - | - | - | 24 |
| ΔEA | 8 | 132.88±1.55 | D'Agostino-Pearson | 0.59 | Yes | 0.25 | Yes | 0.0004 | 23 |
| ΔspaP | 8 | 134.13±0.83 | D'Agostino-Pearson | 0.56 | Yes | 1.00 | Yes | 0.0019 | 23 |
| ΔspaQ | 8 | 132.38±0.74 | D'Agostino-Pearson | 0.54 | Yes | 0.91 | Yes | <0.0001 | 23 |
| ΔspaR | 8 | 132.38±0.92 | D'Agostino-Pearson | 0.73 | Yes | 0.84 | Yes | <0.0001 | 23 |
| ΔinvG | 8 | 137±1.51 | D'Agostino-Pearson | 0.21 | Yes | 0.46 | Yes | 0.1002 | 24 |
| ΔinvA | 8 | 136.63±1.19 | D'Agostino-Pearson | 0.19 | Yes | 0.55 | Yes | 0.1918 | 24 |
| ΔspaS (type 1) | 9 | 135.22±1.64 | D'Agostino-Pearson | 0.01 | No | 0.25 | Yes | 0.3446 | 24 |
| ΔspaS (type 2) | 5 | 131.6±1.67 | Shapiro-Wilk | 0.31 | Yes | 0.18 | Yes | 0.0001 | 23 |

Diameter values are given as average ± standard deviation from n class averages
For all tests the confidence interval is 95%

**Supplementary Table 2:**
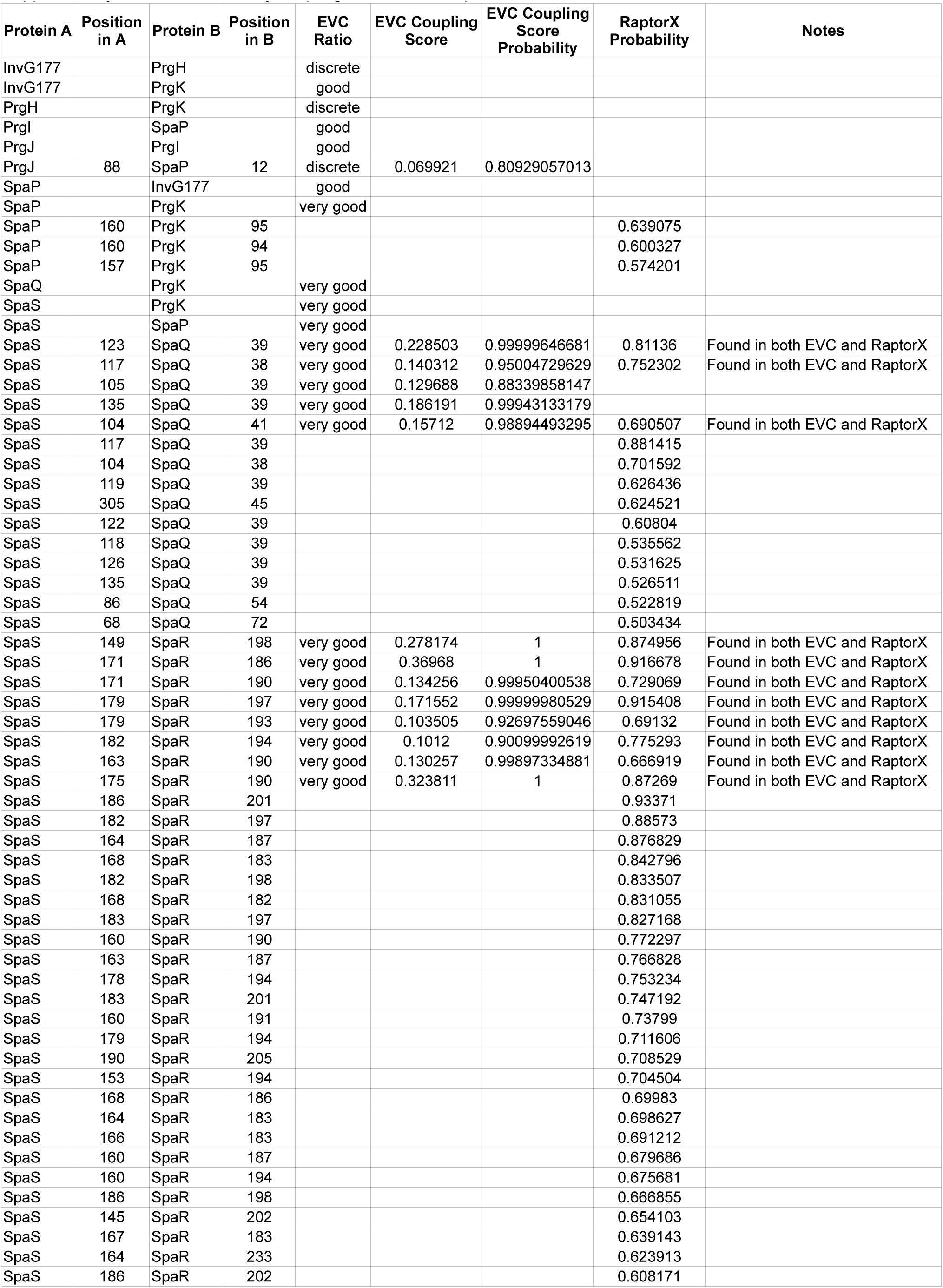
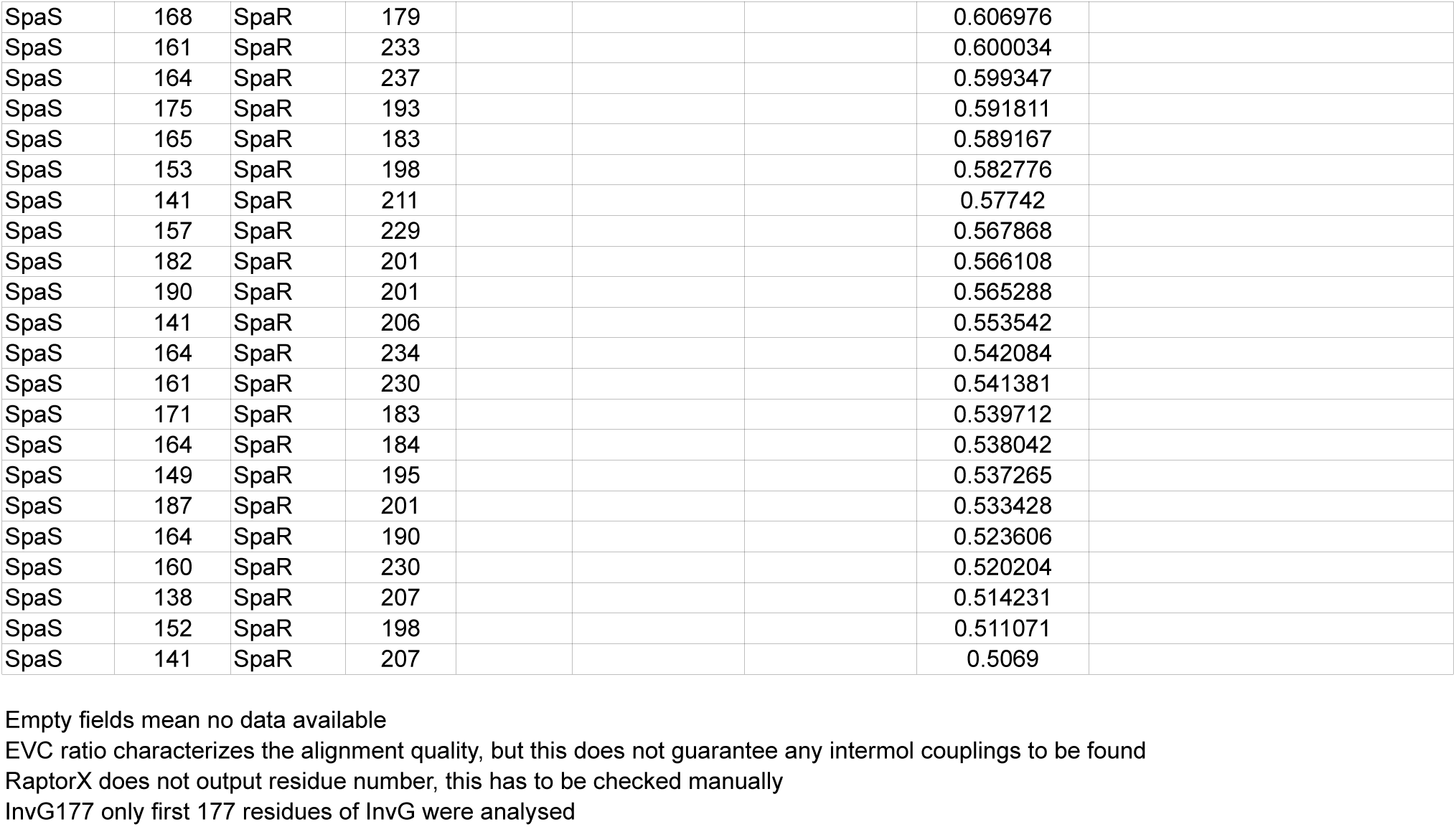
Evolutionary couplings between T3SS proteins.

**Supplementary Table 3:** Cryo-EM data collection summary.

| Sample |  | WT | WT | EA-KO |
| --- | --- | --- | --- | --- |
|  | Buffer | 10 mM Tris-HCl, pH 8.0, 0.5 M NaCl, 5 mM EDTA., 0.1% LDAO |  |  |
|  | Vitrification |  |  |  |
|  | Grid | graphene oxide | carbon | carbon |
| Data collection | Magnification, nominal | 130x | 130x | 130x |
|  | Energy filter slit, eV | not used | 10-15 | 10-15 |
| | Defocus range, $\mu\text{m}$ | 0.7-5.1 | 0.3-5.2 | 0.4-5.1 |
|  | Voltage, kV | 300 | 300 | 300 |
|  | Microscope | Titan Krios | Titan Krios | Titan Krios |
|  | Camera | Gatan K2 | Gatan K2 | Gatan K2 |
|  | Frame exposure time, s | 0.2 | 0.2 | 0.2 |
|  | Dose per frame | 1.24 | 1.26 | 1.1 |
|  | Number of movie frames | 25 | 25 | 25 |
| | Total electron dose, $\text{e}^-/\text{\AA}^2$ | 31 | 31.5 | 27.5 |
| | Pixel size, $\text{\AA}$ | 1.09 | 1.09 | 1.09 |
|  | Number of micrographs<br>(for motioncorrection) | 9352 | 10433 | 8190 |

**Supplementary Table 4:**
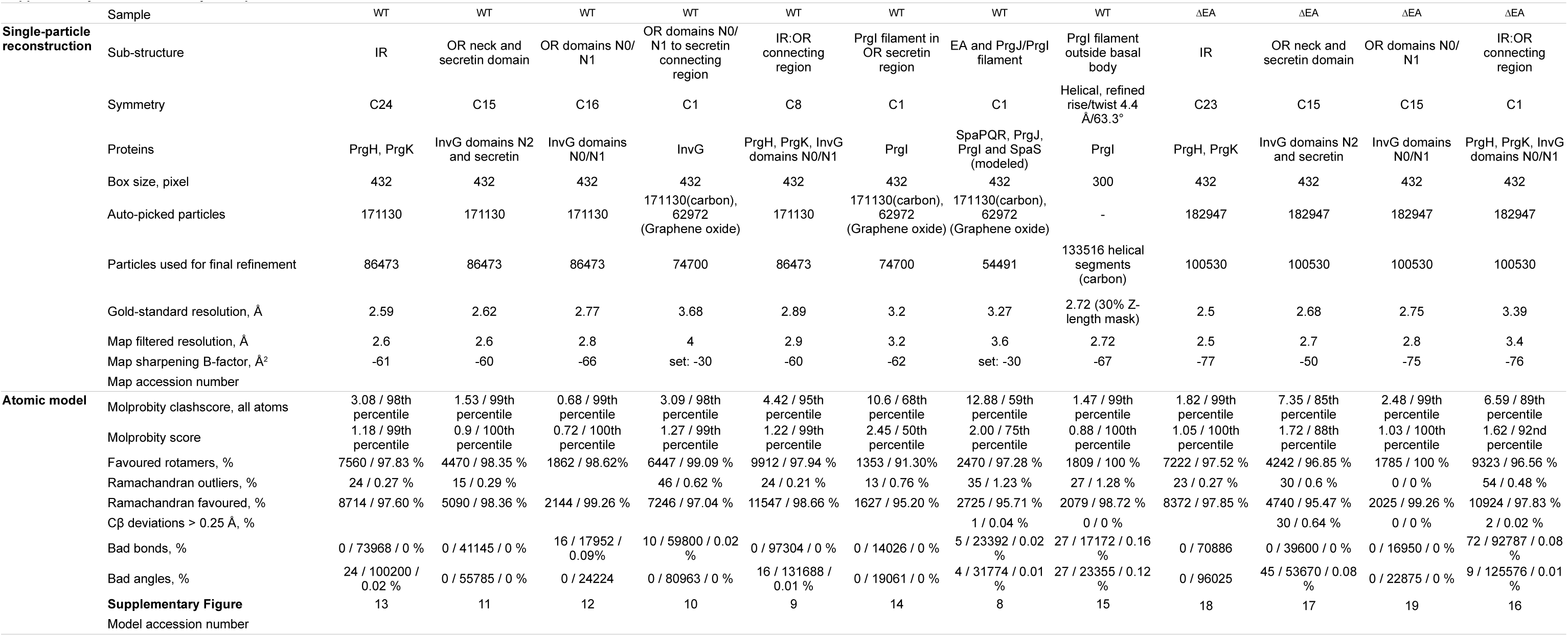
Summary of maps and models.

**Supplementary Table 5:** RMSD between SpaP and SpaQ conformations in EA SpaP.

|  | P1 | P2 | P3 | P4 | P5 |
| --- | --- | --- | --- | --- | --- |
| P1 | - |  |  |  |  |
| P2 | 1.739 | - |  |  |  |
| P3 | 1.489 | 1.478 | - |  |  |
| P4 | 2.871 | 3.446 | 3.104 | - |  |
| P5 | 3.206 | 2.79 | 3.148 | 3.766 | - |

**SpaQ**
|  | Q1 | Q2 | Q3 | Q4 |
| --- | --- | --- | --- | --- |
| Q1 | - |  |  |  |
| Q2 | 1.726 | - |  |  |
| Q3 | 2.165 | 1.32 | - |  |
| Q4 | 2.418 | 2.265 | 2.326 | - |

All values are in Å, calculated in UCSF Chimera after a structure/sequence alignment with MatchMaker

**Supplementary Table 6:**
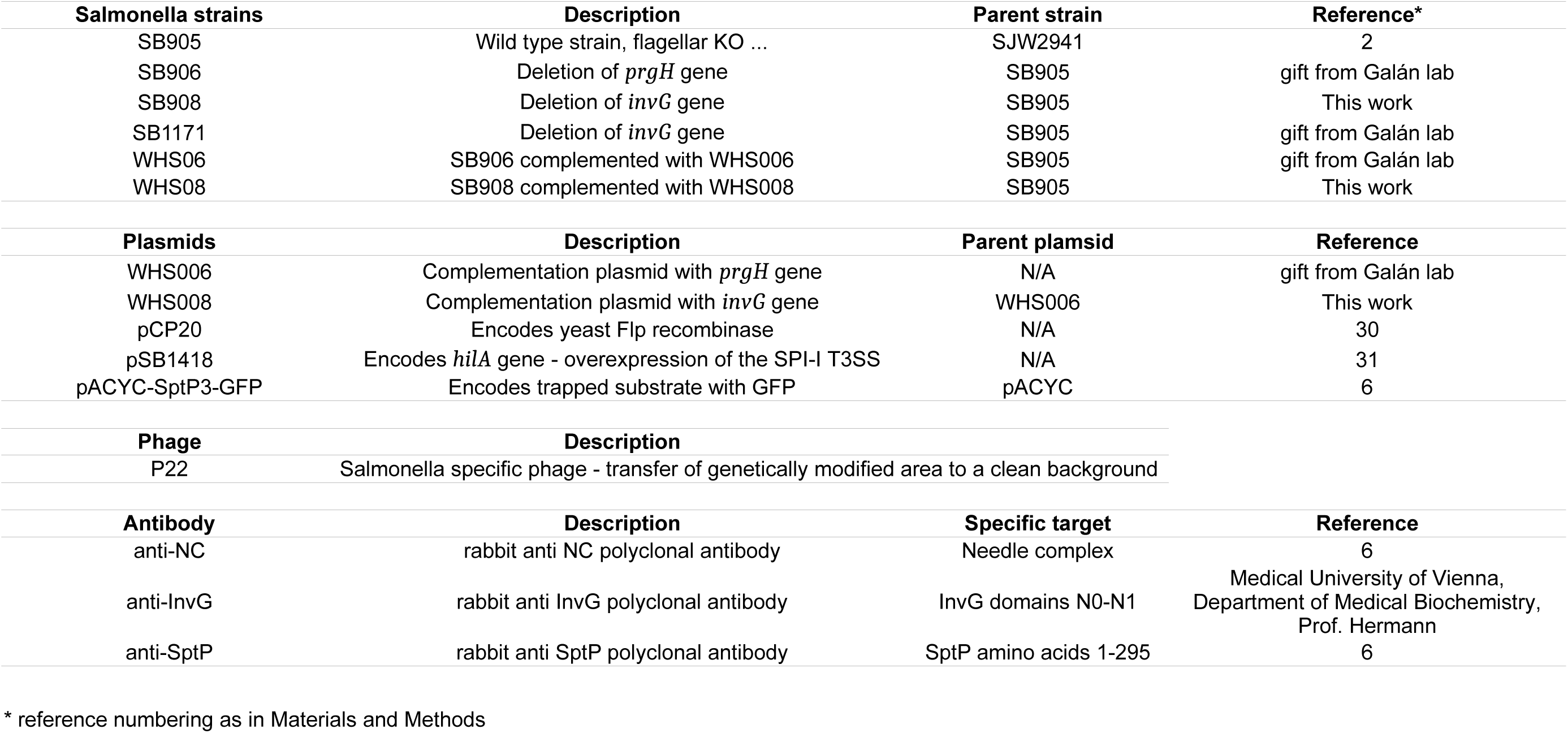
Strains, plasmids and antibodies used in this study.

**Supplementary Table 7:** T3SS protein naming.

| **Protein**<br>**name** | **Unified**<br>**nomenclature** | **UniProt**<br>**ID** | **Flagellar**<br>**homologue** |
| --- | --- | --- | --- |
| InvA | SctV | P0A1I3 | FlhA |
| InvG | SctC | P35672 | - |
| PrgH | SctD | P41783 | FliG |
| PrgI | SctF | P41784 | - |
| PrgJ | SctI | P41785 | - |
| PrgK | SctJ | P41786 | FliF |
| SpaP | SctR | P40700 | FliP |
| SpaQ | SctS | P0A1L7 | FliQ |
| SpaR | SctT | P40701 | FliR |
| SpaS | SctU | P40702 | FlhB |

